# Aging-Associated Microbiota Drives Treg Dysfunction via TNF Signaling

**DOI:** 10.1101/2025.08.08.669389

**Authors:** Jefferson Antônio Leite, Zeynep Ergün, Emmanouil Stylianakis, Rebecca Jasser, Florian Ingelfinger, Natalia Notarberardino Bos, Aysan Poursadegh Zonouzi, Michal Mark, Katlynn Carter, Xinyuan Liu, Nadine Hövelmeyer, Tommy Regen, Manolis Pasparakis, Christian Neumann, Ido Amit, Nir Yosef, Christoph Reinhardt, Ari Waisman

## Abstract

Aging is associated with a chronic, low-grade inflammatory state referred to as inflammaging, which contributes to impaired immune regulation and increased susceptibility to disease. While regulatory T (Treg) cells are key mediators of immune homeostasis, their role in the context of age-related inflammation remains poorly understood. Here we demonstrate that age-related changes in the microbiota promote impaired Treg cell function, resulting in the differentiation of inflammatory T cells. In agreement, we find that aged germ-free (GF) mice exhibited a more balanced immune profile, where the Treg cells are functional and pro-inflammatory mediators are reduced, suggesting that microbial exposure is essential for the establishment of inflammaging. Furthermore, we show that the use of old microbiota in young animals was sufficient to induce pro-inflammatory T cell responses and impaired mucosal Treg cell proliferation, while young microbiota restored Treg cell function in old animals. Mechanistically, we show that exposure to aged microbiota was associated with sustained TNF signaling, elevated oxidative stress, DNA damage, and increased expression of senescence markers such as γH2AX and p16 in Treg cells. These findings uncover a microbiota-TNF-dependent mechanism by which age-associated microbial dysbiosis drives Treg cell dysfunction and promotes immune aging, highlighting the therapeutic potential of microbiota-targeted strategies to restore immune homeostasis in the elderly.

## Introduction

Aging is a natural process that impacts different physiological functions, including immunity, leading to a gradual decline in the ability to mount effective responses to pathogens, tumors, and vaccines. This gradual decline in immune function, referred to as immunosenescence, is characterized by a shift in T cell populations, with a progressive loss of naive T cells and an accumulation of highly differentiated, functionally altered, memory-like subsets. Among these, both effector and regulatory T (Treg) cells undergo significant age-related modifications that contribute to chronic inflammation, or inflammaging, and the pathogenesis of age-associated diseases.^1^

Effector T cells play a crucial role in coordinating immune responses to pathogens and cancer cells, but their characteristics change significantly as the individual ages. A reduction in the naive T cell population, along with an increase of highly differentiated memory-like subsets, results in altered immune responsiveness ^2^. Although these aged T cells can still recognize previously encountered antigens, they show decreased ability to proliferate, limited functional flexibility, and higher levels of markers associated with senescence ^3–5^.

While Treg cells are essential for maintaining immune homeostasis and preventing excessive inflammation, their function during aging remains a topic of ongoing debate. Although there is an increased number of Treg cells in lymphoid and non-lymphoid tissues of older individuals ^6–11^, their ability to suppress immune responses diminishes ^6,10,12,13^. Aged Treg cells display a transcriptional profile that resembles that of effector cells, including increased levels of proinflammatory mediators such as IFN-γ and IL-17A ^10,14–16^. In addition, they exhibit signs of senescence ^10^, including reduced proliferation ^6,10,12,17^. Oxidative stress further compromises their function, contributing to chronic inflammation ^10^. A recent study identified an age-associated subset of Treg cells expressing KLRG1 that accumulates in multiple tissues, displays mitochondrial dysfunction, increased DNA damage, and a pro-inflammatory transcriptional profile, ultimately exhibiting compromised suppressive function *in vivo* despite retaining FOXP3 expression ^18^. However, the molecular mechanisms that lead to the dysregulation of Treg cell function in the elderly are unknown.

Aging corresponds to distinct changes in the gut microbiota, including reduced diversity and shifts in microbial composition^19–21^. Whether these alterations influence immune homeostasis and contribute to inflammaging remains to be determined. Studies have shown that transferring microbiota from old, conventionally housed to young germ-free (GF) mice induces a systemic pro-inflammatory profile characterized by increased activation of CD4⁺ T cells, particularly Th1 and Th17 subsets, highlighting the role of aged microbiota in driving immune activation^22^. Notably, aged murine microbiota was shown to contain increased levels of bacterial taxa linked to inflammation, such as Proteobacteria, and reduced levels of beneficial commensals, like *Akkermansia muciniphila*, which are known to support intestinal barrier integrity and immune regulation^23^.

The relationship between the microbiota, immune function, and aging is increasingly evident. Studies using old GF mice have highlighted the critical role of microbiota in driving age-associated immune dysfunction. Old GF mice exhibit reduced systemic inflammation and maintain better immune homeostasis compared to conventionally raised aged mice. This protection is linked to lower levels of circulating proinflammatory cytokines, reduced intestinal permeability, and preserved macrophage function ^24^. Fecal microbiota transplantation (FMT) from young to old mice reverses several hallmarks of aging, including shifts in the peripheral and brain immune landscape, alongside reduced activation of CD8^+^ T cells ^20,21^.

The decline in short-chain fatty acid (SCFA)-producing bacteria during aging reduces key signals that support the stability and function of Treg cells^20,21,23,25–27^. SCFA-producing bacteria, including *Faecalibacterium*, *Roseburia*, *Eubacterium*, and *Coprococcus*, are critically maintain Treg cell differentiation and function through the production of butyrate, propionate, and acetate, which promote Treg Foxp3 expression and IL-10 production via HDAC inhibition and GPR43 signaling ^28,29^. Simultaneously, increased gut permeability facilitates microbial translocation, exposing Treg cells to persistent inflammatory stimuli like TNF ^24^. In this sense, elevated TNF signaling through TNFR1 has been associated with Treg cells impairment by reducing their proliferation and suppressive function^30^, while TNFR2 activation appears to help sustain Treg cells function^30,31^. Based on these findings, we hypothesize that age-related microbiota changes contribute to Treg cell dysfunction by driving immunosenescence and inflammaging. Moreover, exposure of aged mice to a young microbiota could restore the suppressive function of Treg cells and reduce inflammation, offering a potential strategy to counteract immune dysregulation during aging.

Here, we show that microbial signals are essential drivers of Treg cell dysfunction during aging. Aged GF mice retain functional Treg cells with preserved proliferative capacity and genomic integrity, despite their chronological age. Alternatively, aged specific pathogen free (SPF) mice accumulate dysfunctional Treg cells marked by oxidative stress, DNA damage, senescence, and impaired suppressive function. Microbiota transfer from aged donors is sufficient to induce the dysfunctional Treg phenotype in GF recipients, while young microbiota restores Treg cell stability, demonstrating that dysbiosis is both necessary and sufficient to drive Treg cell dysfunction in aging. Mechanistically, this process is mediated by TNF–TNFR1 signaling induced by the aged microbiota, which promotes oxidative stress and genomic instability in Treg cells. Supporting this mechanism in humans, aged Treg cells from the spleen exhibit a conserved transcriptional signature with NF-κB and STAT1 activation and upregulation of stress and senescence-associated genes, including DDIT4, PPP1R15A, KLF6, and HSPA5, resembling the TNF-driven programs observed in mice.

## Results

### Aging Leads to the Accumulation and Phenotypic Alterations of Regulatory T cells in Mice

Aging is associated with significant shifts in Treg cell homeostasis, but the extent to which these changes impact immune regulation remains unclear. To investigate this, we assessed Treg cell frequency, phenotype, and functional markers in young (2 months) and aged (18 months) C57BL/6J wild-type (WT) mice. Aged mice exhibited a significantly higher frequency of Treg cells in multiple organs, particularly in the spleen, mesenteric lymph nodes (mLNs), and colon **(Fig. 1A)**. In addition, Treg cell number was significantly increased in the spleen of aged mice when compared to young mice. We next evaluated Treg differentiation status by flow cytometry. Among aged Treg cells, there was a marked reduction in the percentage of CD62L expression and a concomitant increase in the percentage expressing CD44, indicating a shift toward an effector memory-like phenotype **(Fig. 1B)**. To further dissect the heterogeneity of Treg cells in aging, we performed high-dimensional flow cytometry analysis of splenocytes from young and old mice. Unsupervised clustering revealed distinct Treg cell subsets, with aged mice displaying an enrichment of certain clusters, including KLRG1^+^, GATA-3^+^ST2^+^ and PD-1^+^ Treg cells **(Fig. 1C)**. We next examined specific Treg cell subsets associated with immune regulation and inflammatory responses. Colonic Helios+ and RORγt⁺ Treg cell frequencies remained unchanged with aging **(Fig. 1D)**. Since KLRG1 is associated with terminal differentiation^32^, we next assessed its expression in aged Treg cells. Aged mice exhibited a significantly higher frequency of KLRG1^+^ Treg cells in the spleen **(Fig. 1E)** and colon (not shown), supporting the hypothesis that aging is associated with the accumulation of more terminally differentiated Treg cells. In contrast, ST2^+^ Treg cells, involved in maintaining tissue homeostasis, remained unchanged between young and aged mice **(Fig. 1F)**. This suggests that aging affects specific subsets of Treg cells.

**Figure 1:**
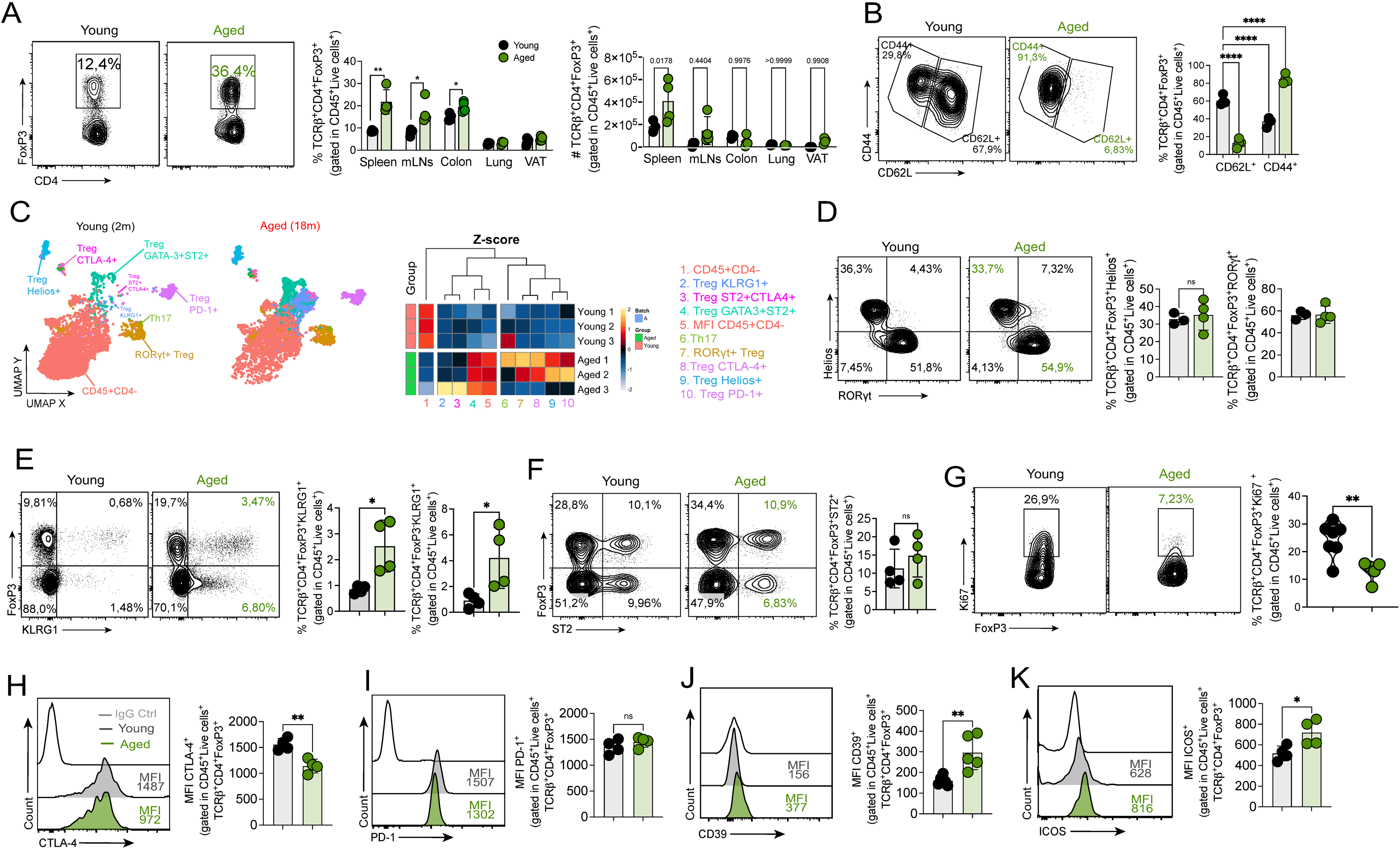
Aging Remodels Treg Cell phenotype, function, and stability across tissues. (**A**) Representative dot plots showing CD4⁺Foxp3⁺ Treg cell frequencies in the spleen of young (2-month-old) and aged (18-month-old) C57BL/6 wild-type mice. Adjacent bar graphs quantify Treg cell frequencies (%) and absolute Treg cell numbers across the spleen, mesenteric lymph nodes (mLNs), colon, lung, and VAT. Data represent mean ± SEM. Statistical significance was assessed using unpaired t-test. (**B**) Representative dot plots of CD44 and CD62L expression in CD4⁺Foxp3⁺ Treg cells from young and aged mice. The adjacent bar graph quantifies CD44⁺CD62L⁻ effector-like Treg cells (%). Data shown as mean ± SEM. (**C**) t-SNE plots from high-dimensional flow cytometry of CD4⁺ T cells in young and aged mice, identifying distinct clusters. The adjacent heatmap displays Z-score normalized frequencies of major clusters, including Treg cells and effector T cells. (**D**) Representative dot plots of Helios and Rorgt expression in Foxp3+ CD4⁺ T cells from young and aged mice. Adjacent bar graphs quantify Helios⁺ Treg cells (%) and RORγt⁺ Treg cells (%). Data are mean ± SEM. (**E**) Representative dot plots of KLRG1 and Foxp3 expression within CD4⁺ from the spleen of young and aged mice. The adjacent bar graph quantifies KLRG1⁺Foxp3⁺ Treg cells and KLRG1⁺Foxp3^-^CD4^+^ T cells (%). (**F**) Representative FACS dot plots of ST2 and Foxp3 expression within CD4⁺ T cells from young and aged mice. The adjacent bar graph quantifies ST2⁺ Treg cells (%). (**G**) Representative histograms of Ki67 expression in CD4⁺Foxp3⁺ Treg cells from young and aged mice. The adjacent bar graph quantifies Ki67 MFI.. (**H**) Representative histograms of CTLA-4 expression in CD4⁺Foxp3⁺ Treg cells rom young and aged mice. The adjacent bar graph quantifies CTLA-4 MFI. Data shown as mean ± SEM. (I) Representative histograms of PD-1 expression in CD4⁺Foxp3⁺ Treg cells from young and aged mice. The adjacent bar graph quantifies PD-1 MFI. (J) Representative histograms of CD39 expression in CD4⁺Foxp3⁺ Treg cells from young and aged mice. The adjacent bar graph quantifies CD39 MFI. Data shown as mean ± SEM. (K) Representative histograms of ICOS expression in CD4⁺Foxp3⁺ Treg cells from young and aged mice. The adjacent bar graph quantifies ICOS MFI. Data represent mean ± SEM. Statistical analysis for panels A–K was performed using unpaired t-test. *p < 0.05; **p < 0.01. Data shown are representative of two to three independent experiments (n = 4–5 mice per group) and presented as mean ± SEM. Each symbol represents one mouse. Representative of three independent experiments with n = 4 mice per group.

To determine whether aging impacts Treg cell renewal, we analyzed Ki67 expression, a marker of cell proliferation. We found that significantly fewer aged Treg cells expressed Ki67, indicating diminished proliferative capacity **(Fig. 1G)**. We next assessed the expression of key immunoregulatory molecules that contribute to Treg cell function. Among aged Treg cells, a significant reduction in CTLA-4 expression was observed **(Fig. 1H)**, while PD-1 expression was unchanged **(Fig 1I)**, and CD39 **(Fig. 1J)** and ICOS **(Fig. 1K)** were significantly upregulated.

Together, these results indicate that aging leads to an increased frequency of Treg cells, and that these cells undergo impaired proliferation.

### Aged Treg Cells Exhibit Impaired Function and Fail to Suppress Intestinal Inflammation

We next investigated how phenotypic alterations in aged Treg cells impact their function in steady-state conditions and in a model of T cell-driven colitis. To determine whether aging influences the baseline activation and cytokine profile of T cells, we first analyzed the frequency of IFN-γ^+^ and IL-17A^+^ CD4^+^ and CD8^+^ T cells in young and aged mice under steady-state conditions. Aged mice exhibited an increase in IFN-γ^+^ CD4^+^ and CD8^+^ T cells, while the frequency of IL-17A^+^ CD4^+^ T and CD8+ T cells remained unchanged compared to young mice, particularly in lymphoid organs such as the spleen (SP), mLNs, and the small and large intestinal lamina propria (SI-LP and LI-LP) **(Fig. 2A, B, C)**.

**Figure 2:**
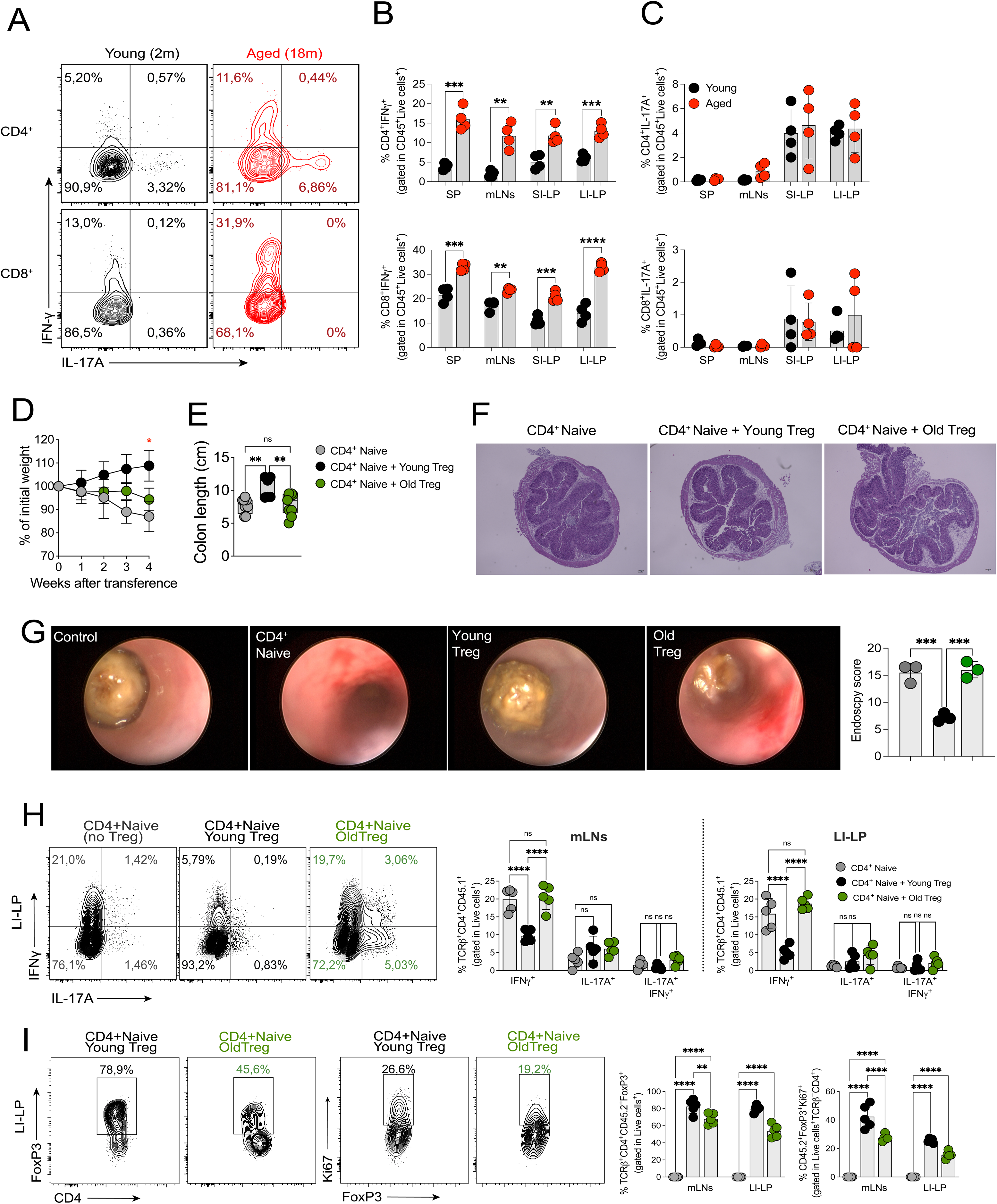
Aged Treg cells fail to Suppress Intestinal Inflammation and Pro-Inflammatory T Cell Responses. (**A**) Representative dot plots showing frequencies of IFN-γ⁺ and IL-17A⁺ CD4⁺ T cells (top) and CD8⁺ T cells (bottom) in spleen (SP) and mesenteric lymph nodes (mLNs) from young and aged mice under steady-state conditions. (B, C) Quantification of IFN-γ⁺ CD4⁺ T cells (top left), IL-17A⁺ CD4⁺ T cells (top right), IFN-γ⁺ CD8⁺ T cells (bottom left), and IL-17A⁺ CD8⁺ T cells (bottom right) in spleen SP, mLNs, small intestinal lamina propria (SI-LP), and large intestinal lamina propria (LI-LP) of young and aged mice. (D) Body weight curve over time (4 weeks) in Rag1⁻/⁻ mice transferred with CD4⁺ naïve T cells alone, or co-transferred with young or aged Treg cells. (E) The bar graph shows colon length at the experimental endpoint. (F) Representative H&E-stained colon sections from control mice, mice receiving CD4⁺ naïve T cells alone (left), or co-transferred with young (middle) or aged Treg cells (right). (G) Representative colonoscopic images showing colon inflammation in mice transferred with CD4⁺ naïve T cells alone, or co-transferred with young or aged Treg cells. Control mice (no transfer) are also shown. The adjacent graph quantifies endoscopic scores. (H) Representative dot plots showing IFN-γ and IL-17A expression in CD4⁺ T cells from mLNs and LI-LP of mice transferred with CD4⁺ naïve T cells alone, or co-transferred with young or aged Treg cells. Quantification of IFN-γ⁺ and IL-17A⁺ CD4⁺ T cells in mLNs (left) and LI-LP (right). (I) Representative dot plots showing frequencies of Foxp3⁺ Treg cells in mLNs and LI-LP of mice co-transferred with young or aged Treg cells and quantification of Ki67⁺ proliferating Treg cells (Foxp3⁺Ki67⁺) in mLNs and LI-LP of mice co-transferred with young or aged Treg cells. Data shown are representative of two to three independent experiments (n = 4–5 mice per group) and presented as mean ± SEM. Statistical analysis was performed using ANOVA with post-hoc test or unpaired t-test. *p < 0.05; **p < 0.01. Each symbol represents one mouse.

To evaluate whether aged Treg cells retain their suppressive function *in vivo*, we used a well-established model of T cell-driven colitis. Rag1-deficient mice received naïve CD45.1^+^ CD4^+^ T cells alone or in combination with CD45.2^+^ Treg cells from young or aged mice. Body weight loss was monitored as an indicator of disease severity. Mice receiving aged Treg cells exhibited significantly greater weight loss compared to those receiving young Treg cells, suggesting that aged Treg cells are less effective at controlling colitis progression **(Fig. 2D)**. Likewise, colon length, a marker of colitis severity, was significantly shorter in mice receiving aged Treg cells compared to those receiving young Treg cells **(Fig. 2E)**, suggesting increased tissue damage and inflammation in the presence of aged Treg cells. Histological analysis using H&E staining of colonic tissue and mini-endoscopy analysis further confirmed that mice receiving aged Treg cells exhibited increased infiltration of immune cells compared to those receiving young Treg cells **(Fig. 2F, G)**.

To directly assess the inflammatory response in the intestine, we quantified IFN-γ^+^ and IL-17A^+^ CD4^+^ T cells in the lamina propria. Mice receiving young Treg cells exhibited a significant reduction in these populations LI-LP, whereas mice receiving aged Treg cells displayed persistent IFN-γ^+^ and IL-17A^+^ CD4^+^ T cell accumulation **(Fig. 2H)**. To investigate potential mechanisms underlying the defective function of aged Treg cells, we assessed their persistence *in vivo*. The frequency of Foxp3^+^ Treg cells in the mLNs and LI-LP was significantly lower in mice receiving aged Treg cells, indicating a diminished capacity of aged Treg cells to expand and persist in an inflammatory environment **(Fig. 2I)**. Supporting this data, a significantly lower percentage of aged Treg cells expressed Ki67^+^ **(Fig. 2I)**, suggesting a reduced proliferative capacity, which may contribute to their inability to control inflammation.

Together, these results indicate that aging leads to a functional decline in Treg cells, reducing their ability to maintain intestinal immune homeostasis and suppress inflammatory T cell responses. The impaired expansion and persistence of aged Treg cells, along with their failure to control IFN-γ^+^ and IL-17A^+^ effector T cells, coincides with increased intestinal inflammation and disease severity.

### Microbiota Transfer Modulates T Cell Responses and Gut Microbiome Composition in Aging

Aging is associated with significant alterations in the gut microbiota that have been implicated in immune dysfunction^20,21,23,24^. Our findings in Figs. 1 and 2 demonstrated that aged mice accumulate Treg cells with impaired suppressive capacity, reduced proliferative potential, which fail to control excessive inflammation. Given the established role of microbiota in shaping immune responses and Treg cell-mediated tolerance, we investigated whether restoring a youthful microbiota could rescue Treg cell function and mitigate inflammatory signatures in aged mice. To this end, we treated young and aged mice with a broad-spectrum antibiotic cocktail (ABX) to deplete their endogenous microbiota, and then performed fecal microbiota transplantation (FMT) using material from either young or aged donor mice. The frequency of Foxp3⁺ Treg cells in the spleen **(Fig. 3A upper panel)**, mLNs, and LI-LP (data not shown) did not change in any of the experimental groups. Young mice that received aged microbiota (blue squares) did not show an increase in Treg cell frequency compared to young mice, and aged mice that received young microbiota (red squares) also maintained similar frequencies of Treg cells compared to young ones **(Fig. 3A, upper panel)**. In contrast, the expression of Ki67 in Treg cells was affected by the microbiota composition. Young mice that received aged microbiota showed a reduction in the proportion of Ki67⁺ Treg cells, whereas aged mice that received young microbiota showed increased proportion of Ki67⁺ Treg cells **(Fig 3A, lower panel)**, suggesting improved proliferative capacity. These findings suggest that although the total number of Treg cells is not regulated by the microbiota, microbial signals influence their proliferation, and potentially, their suppressive function. Further, inflammatory signals derived from the aged microbiota likely compromise Treg cell stability and function.

**Figure 3:**
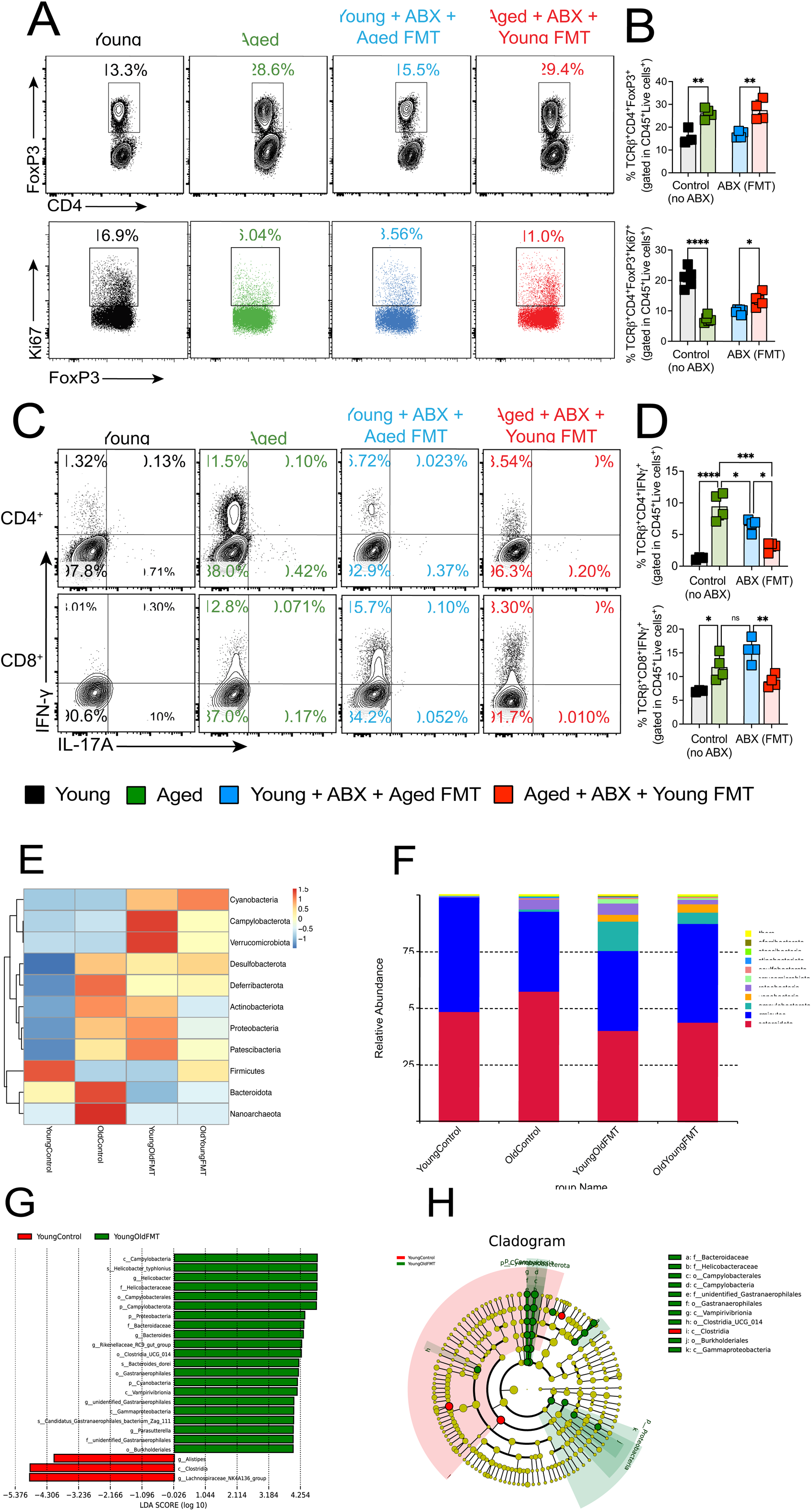
Microbiota transfer modulates Treg homeostasis and inflammatory T cell responses. (A) Representative dot plots showing CD4⁺Foxp3⁺ Treg frequencies in the spleen of aged mice, young mice treated with antibiotics (ABX) and transplanted with aged microbiota (Aged FMT), and aged mice treated with ABX and transplanted with young microbiota (Young FMT). (B) Adjacent bar graphs quantify Treg frequencies (%) and Ki67⁺ proliferating Treg cells (%). (C) Representative dot plots showing frequencies of IFN-γ⁺ CD4⁺ T cells and IFN-γ⁺ CD8⁺ T cells in the spleen across the same experimental groups as panel A. (D) Adjacent bar graphs quantify the percentages of IFN-γ⁺ CD4⁺ T cells (top) and IFN-γ⁺ CD8⁺ T cells (bottom) (E) Heatmap of Z-score normalized microbial taxa abundances in fecal samples from young control and young + aged FMT groups, highlighting shifts in microbial composition. (F) Bar plots showing relative abundances of bacterial phyla across young control, young + aged FMT, and aged + young FMT groups. (G) Linear discriminant analysis effect size (LEfSe) plot identifying bacterial taxa significantly enriched in young control (green) versus aged FMT (red) groups. (H) Cladogram showing phylogenetic distribution of bacterial taxa that differ between young control and young + aged FMT groups. Taxa enriched in young controls are marked in green, while those enriched in aged FMT recipients are in red. Data shown are representative of two to three independent experiments (n = 4–5 mice per group) and presented as mean ± SEM. Statistical analysis was performed using ANOVA with post-hoc test or unpaired t-test. *p < 0.05; **p < 0.01. Each symbol represents one mouse.

Since Treg cells from young mice displayed reduced proliferative capacity after exposure to aged microbiota (**Fig. 3A, B**), we next evaluated whether these microbiota-induced changes also extended to effector T cell responses. Aged mice that received young microbiota showed a marked reduction in the frequency of IFN-γ⁺ CD4⁺ and CD8⁺ T cells **(Fig. 3C)**, indicating that reconstitution with a youthful microbial community can attenuate age-associated inflammatory T cell activity. In contrast, young mice colonized with aged microbiota exhibited increased proportions of IFN-γ-producing T cells **(Fig. 3C, D)**, demonstrating that the aged microbiota alone is sufficient to enhance pro-inflammatory T cell responses.

Since we observed that the aged microbiota influences both Treg cell homeostasis and inflammatory T cell responses, we next sought to determine whether these immune alterations were associated with specific microbial changes. To address this, we performed 16S rRNA sequencing to compare the microbiota composition across the experimental groups. Analysis of microbial abundance at the phylum level showed that mice harboring aged microbiota, including aged mice and young mice that received aged microbiota, exhibited an increased relative abundance of Proteobacteria and Patescibacteria, compared to young controls and aged mice colonized with young microbiota **(Fig. 3E)**. These shifts were corroborated by the relative abundance profiles, which further confirmed the expansion of Proteobacteria and the depletion of Firmicutes and Bacteroidota in mice exposed to aged microbiota **(Fig. 3F)**.

To further characterize the microbial differences, linear discriminant analysis (LDA) identified key bacterial taxa that were differentially enriched among groups. Bacteroidaceae, Helicobacteraceae, Campylobacterales, and Gammaproteobacteria were significantly enriched in young mice receiving aged microbiota **(Fig. 3G)**. In contrast, taxa such as Alistipes, Clostridia, and Lachnospiraceae were enriched in young control mice. In addition, the cladogram shows that the transfer of aged microbiota drives the expansion of specific lineages, particularly within Proteobacteria, including members of Helicobacteraceae and Gammaproteobacteria, while beneficial commensal groups, such as Firmicutes (e.g., Lachnospiraceae) and Bacteroidota, are reduced or underrepresented in recipients of aged microbiota **(Fig. 3H)**.

These findings suggest that aging-associated microbiota disrupts Treg cell proliferation capacity and promotes a pro-inflammatory immune profile. The reversal of these changes by young microbiota transplantation suggests that microbial dysbiosis is a key factor in driving Treg cell dysfunction and immune aging.

### Young microbiota rejuvenates old Treg cells

Our previous findings suggest that an aged microbiota composition affects Treg cell homeostasis and promotes inflammatory T cell responses. To investigate whether this process could be reversed, we cohoused (CH) aged mice with young mice for four weeks and compared their immune profiles to aged mice maintained in single housing (SH). Flow cytometry analysis revealed that aged SH mice exhibited markedly increased frequencies of IFN-γ⁺ and IL-17A⁺ CD4⁺ and CD8⁺ T cells compared to young controls, indicating a heightened inflammatory state (**Fig 4A**). Interestingly, cohousing significantly reduced the frequency of these pro-inflammatory subsets, suggesting that microbiota transfer from young mice mitigates age-associated inflammation **(Fig. 4A).**

**Figure 4:**
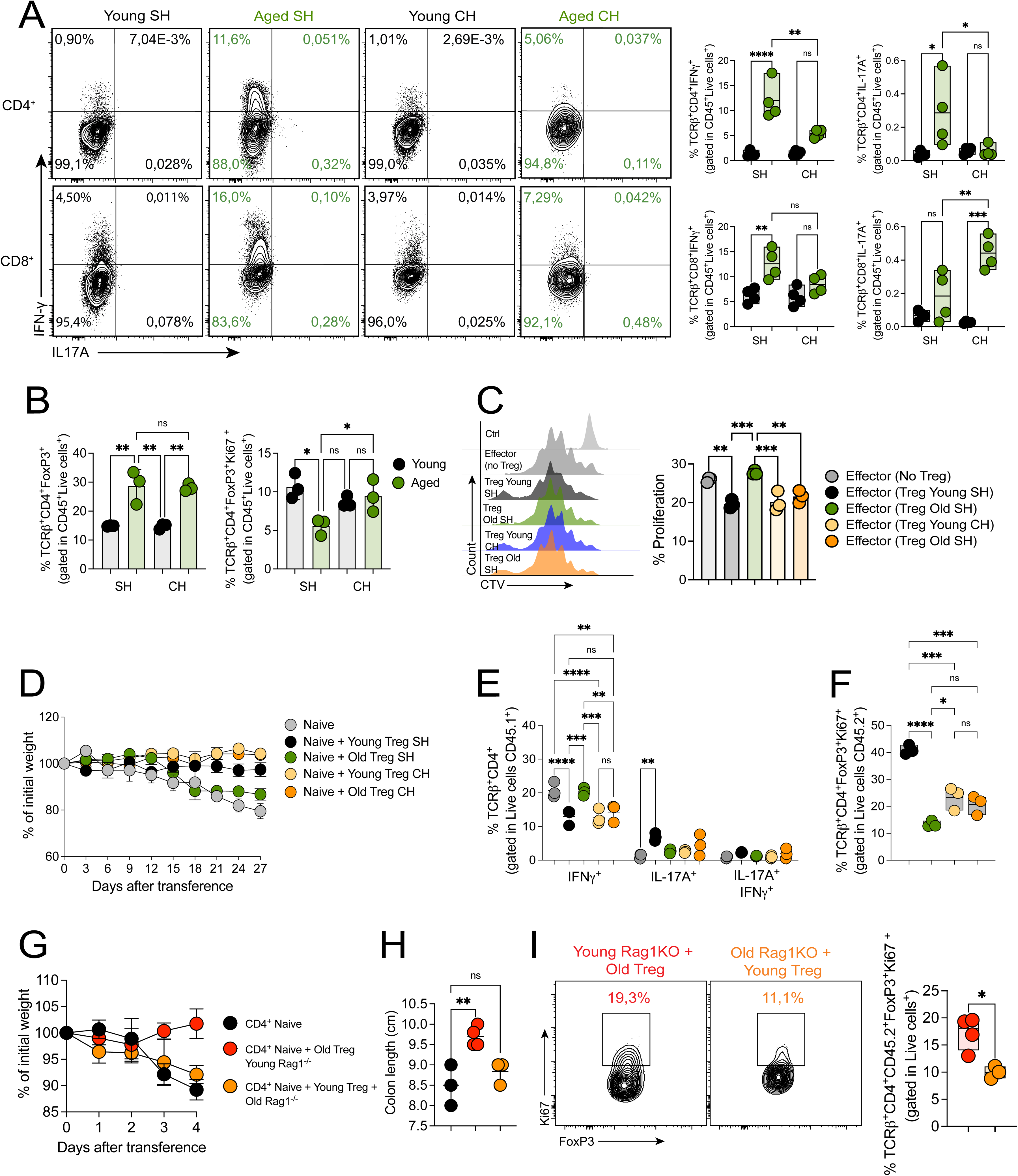
Co-housing with young mice restores Treg cell function and stability in aged mice. (A) Representative dot plots showing IFN-γ and IL-17A expression in CD4⁺ and CD8⁺ T cells from the spleen of aged single-housed (SH) and co-housed (CH) mice. Adjacent bar graphs quantify the frequencies of IFN-γ⁺ and IL-17A⁺ T cells within CD4⁺ and CD8⁺ populations. Data represent mean ± SEM. (B) Quantification of Treg cell frequencies (% CD4⁺Foxp3⁺) and proliferating Treg cells (% Foxp3⁺Ki67⁺) in the spleen of young and aged mice kept in SH or CH with young mice for 4 weeks. (**C**) *In vitro* suppression assay evaluating the ability of Treg cells from aged single-housed (SH) or CH mice to suppress CD4⁺ T cell proliferation. Effector CD4⁺ T cells were labeled with CellTrace Violet (CTV) and cultured with or without Treg cells. Representative CTV dilution histograms are shown for each condition. The adjacent bar graph quantifies the proliferation index of effector T cells across conditions. (**D**) Body weight curves of Rag1⁻/⁻ mice transferred with CD4⁺ naïve T cells alone or co-transferred with young or aged Treg cells from SH or CH conditions. (E) Quantification of IFN-γ⁺ and IL-17A⁺ CD4⁺ T cells in the mesenteric lymph nodes (mLNs) of recipient mice from panel D. (F) Quantification of Ki67⁺ proliferating Treg cells in the mLNs of recipient mice from panel D (G) Body weight curves and (H) colon length of Rag1⁻/⁻ mice preconditioned with young or aged Treg cells and subsequently transferred with CD4⁺ naïve T cells. (I) Representative dot plots showing Ki67 expression in Foxp3⁺ Treg cells from Rag1⁻/⁻ recipient mice preconditioned with young or aged Treg cells. The adjacent bar graph quantifies Ki67⁺ Treg cells (%). Data shown are representative of two to three independent experiments (n = 3–5 mice per group) and presented as mean ± SEM. Statistical analysis was performed using ANOVA with post-hoc test or unpaired t-test. *p < 0.05; **p < 0.01. Each symbol represents one mouse.

We next assessed whether microbiota remodeling also impacts Treg cell homeostasis. While the overall frequency of Treg cells remained unchanged between SH and CH aged mice, the frequency of Treg cells expressing Ki67 was significantly increased in CH mice, indicating improved proliferative capacity **(Fig. 4B).** To evaluate whether this proliferative improvement was accompanied by functional restoration, we performed an *in vitro* suppression assay. Indeed, Treg cells from CH aged mice exhibited enhanced suppressive capacity, as shown by reduced proliferation of effector T cells compared to Treg cells from SH aged mice, demonstrating that exposure to a young microbiota restores Treg-mediated suppression **(Fig. 4C)**.

To investigate whether these effects translate to an *in vivo* setting, we performed Treg cell transfer experiments in a T-cell transfer colitis model in Rag1^-/-^ mice. To this end, young and aged mice were left in SH versus CH conditions for four weeks and after this time, Treg cells from both SH and CH groups were isolated and co-transferred with naïve CD4 T cells into Rag1^-/-^ animals. Mice that received aged Treg cells from SH mice developed more severe disease, as evidenced by greater weight loss **(Fig. 4D)** and increased IFN-γ and IL-17A production in the colon **(Fig. 4E)**. In contrast, the transfer of Treg cells from CH aged mice led to improved disease outcomes, with reduced inflammation-associated cytokines. Additionally, aged Treg cells from CH mice exhibited increased proliferation, although it was not statistically significant **(Fig. 4F).**

We next investigated whether the aged environment drives Treg cell dysfunction and whether a young environment can restore Treg cell function. To this end, aged Treg cells, which exhibited impaired suppressive capacity, were first transferred into young Rag1⁻^/^⁻ mice and maintained in this environment for four weeks before co-transfer with naïve CD4 T cells **(Fig. 4G, H)**. In addition, young Treg cells, which retain suppressive function, were transferred into aged Rag1⁻^/^⁻ mice and kept in this aged environment for four weeks before naïve CD4 T cell transfer (red group). Following naïve T cell transfer, weight monitoring revealed that young Rag1⁻^/^⁻ mice preconditioned with aged Treg cells exhibited less weight loss and reduced colitis severity. In contrast, aged Rag1⁻^/^⁻ mice preconditioned with young Treg cells developed more severe disease, as indicated by greater weight loss **(Fig. 4G)**. Furthermore, analysis of Ki67 expression in Treg cells revealed that young Rag1⁻^/^⁻ mice preconditioned with aged Treg cells exhibited higher Treg proliferation compared to aged Rag1⁻^/^⁻ mice preconditioned with young Treg cells **(Fig. 4I)**. These findings support the idea that exposure to a youthful microbiota can restore aged Treg cell function. This effect is likely mediated by microbial signals that enhance Treg proliferation and suppressive capacity, as indicated by increased Ki67 expression and improved control of inflammation. Thus, a young microbiota helps maintain Treg stability and immune homeostasis during aging.

### Aged Microbiota Drives Pro-Inflammatory T Cell Differentiation and Impairs Treg Cell Homeostasis

Our previous findings suggest that aged microbiota impairs Treg cell proliferation to promote a pro-inflammatory immune environment. To investigate how microbiota composition influences T cell differentiation and cytokine production, we reanalyzed publicly available single-cell RNA sequencing (scRNA-seq) data from Kawamoto et al. (2023)^33^, comparing T cell populations under SPF and GF conditions in the lamina propria of young and aged mice.

UMAP clustering revealed distinct T cell populations across GF and SPF conditions, highlighting the impact of microbiota on immune differentiation **(Fig. 5A)**. Proportional analysis showed that aged SPF mice exhibited a marked increase in inflammatory subsets, including effector CD8^+^ T cells, compared to young SPF mice **(Fig. 5B)**. In contrast, aged GF mice retained a more balanced T cell profile, suggesting that exposure to an aged microbiota environment drives pro-inflammatory T cell differentiation. In addition, effector T cells from aged SPF mice upregulated exhaustion markers such as Lag3, Tox, Tigit and Tcf7, while aged GF mice had a less activated transcriptional profile **(Fig. 5C)**. Similarly, analysis of Treg cell-associated genes revealed that aged SPF mice displayed alterations in key regulatory markers, including Foxp3, ICOS, and IL-10 **(Fig. 5D).**

**Figure 5:**
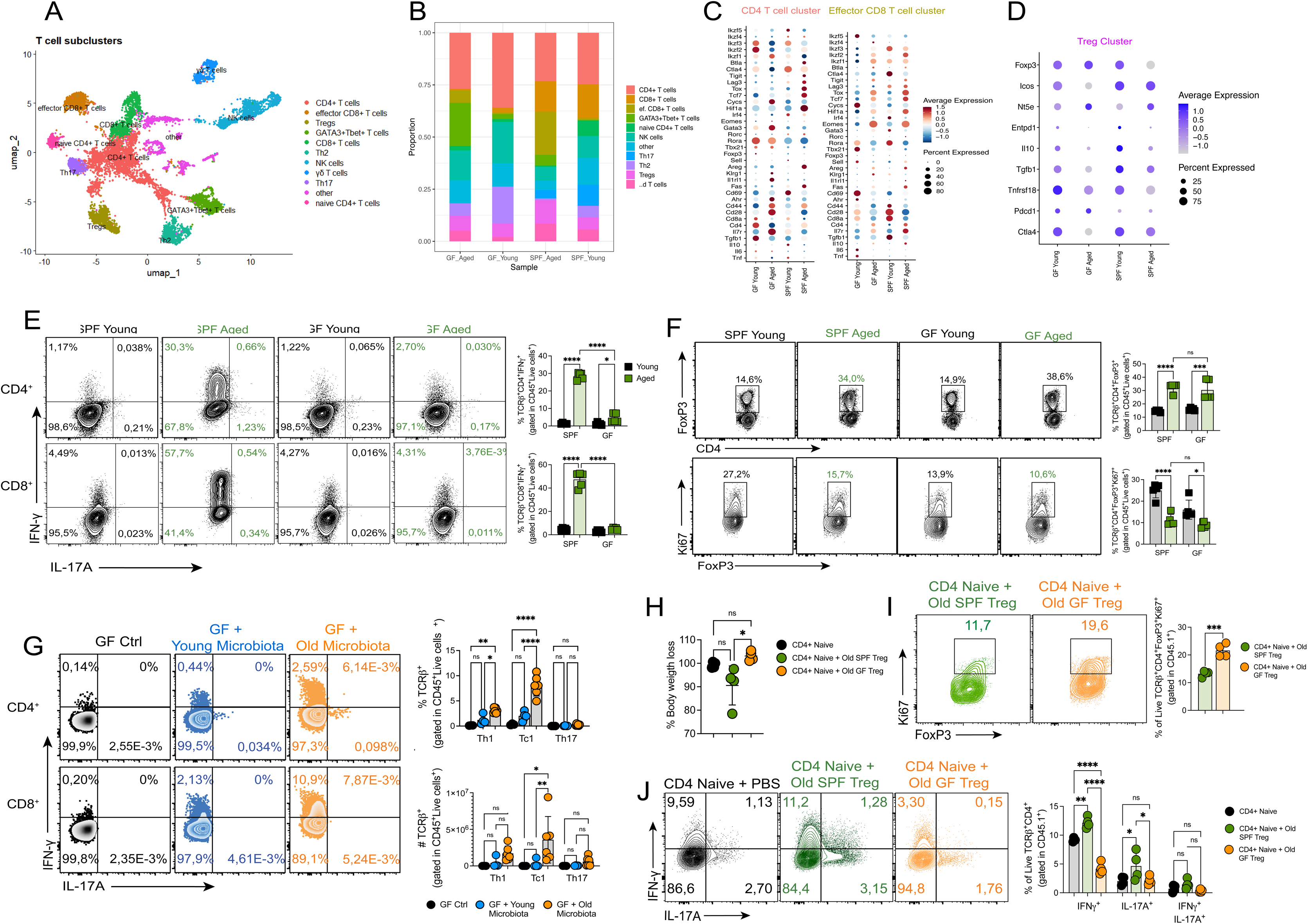
Microbiota modulates Treg cell homeostasis and inflammatory T cell responses during aging. (A) UMAP plots of CD4⁺ and CD8⁺ T cell clusters from germ-free (GF) and specific pathogen-free (SPF) young and aged mice based on single-cell RNA sequencing data. Major subsets, including Treg cells, Th1, Th17, and effector CD8⁺ T cells, are indicated. (B) Proportional abundance of T cell subsets across GF-aged, GF-young, SPF-aged, and SPF-young groups, based on clustering from scRNA-seq. (C) Dot plots showing expression of signature genes across CD4⁺ T cell clusters (left), effector CD8⁺ T cell clusters (middle), and (D) Treg clusters (right), highlighting differences in gene expression patterns between GF and SPF mice. (E) Representative dot plots showing IFN-γ⁺ CD4⁺ T cells and IFN-γ⁺ CD8⁺ T cells from SPF-aged and GF-aged mice. Adjacent bar graphs quantify IFN-γ⁺ T cell frequencies in these groups. (F) Representative dot plots showing Foxp3⁺ Treg frequencies in SPF-aged and GF-aged mice. Adjacent bar graphs quantify Treg frequencies and Ki67⁺ proliferating Treg cells in these groups. Data represent mean ± SEM. Unpaired t-test was used. (G) Representative dot plots showing IFN-γ⁺ CD4⁺ T cells and IFN-γ⁺ CD8⁺ T cells in GF mice colonized with either young microbiota or aged microbiota. Adjacent scatter plots quantify these populations.(H) Body weight loss of Rag1⁻/⁻ recipient mice transferred with CD4⁺ naïve T cells alone, or co-transferred with old Treg cells from either SPF-aged or GF-aged mice. Weight monitoring was performed over time post-transfer. (I) Quantification of Ki67⁺ proliferating Foxp3⁺ Treg cells in the spleen of recipient mice from panel H (J) Quantification of IFN-γ⁺ and IL-17A⁺ CD4⁺ T cells in the large intestinal lamina propria (LI-LP) of recipient mice from panel H. Data shown are representative of two to three independent experiments (n = 3–7 mice per group) and presented as mean ± SEM. Statistical analysis was performed using ANOVA with post-hoc test or unpaired t-test. *p < 0.05; **p < 0.01. Each symbol represents one mouse.

Next, we performed flow cytometry analysis to compare T cell subsets between young and aged SPF and GF mice. We observed a significant reduction in IFN-γ⁺ CD4⁺ and CD8⁺ T cells in aged GF when compared to aged SPF mice, supporting the conclusion that the microbiota is a key driver of the pro-inflammatory phenotype associated with aging **(Fig. 5E)**. In addition, aged GF mice exhibited a similar frequency of Foxp3⁺ Treg cells to aged SPF mice, when compared to young SPF and GF controls. However, these Treg cells displayed reduced Ki67 expression **(Fig. 5F).**

To directly assess whether aged microbiota influences T cell differentiation, we colonized young GF mice with microbiota from either young or aged SPF donors. Interestingly, GF mice receiving aged microbiota developed an increased frequency of IFN-γ⁺ and IL-17A⁺ T cells, while GF mice colonized with young microbiota maintained a more homeostatic immune profile **(Fig. 5G)**. In addition, GF mice that received young microbiota displayed a significantly increased percentage of Ki67⁺ Tregs when compared to those colonized with aged microbiota. This effect was most evident in mucosal sites, where Treg proliferation was significantly higher in GF mice colonized with young microbiota, particularly in mLNs and LI-LP **(Sup.** Fig. 1A, B).

To functionally evaluate whether aged microbiota affects the suppressive capacity of Treg cells, we utilized a T cell transfer colitis model using CD4⁺ naïve T cells co-transferred with Treg cells isolated from aged SPF or GF mice. Interestingly, mice that received Treg cells from aged GF donors exhibited less weight loss during colitis development compared to those receiving aged SPF Treg cells, suggesting a preserved suppressive function in the absence of microbiota-derived inflammatory stimuli **(Fig. 5H)**. This effect was accompanied by a higher proportion of Ki67⁺ Treg cells and a lower frequency of IFN-γ⁺ and IL-17A^+^ CD4⁺ T cells in the LI-LP **(Fig. 5I, J**). These findings indicate that while an aged microbiota promotes Treg dysfunction and inflammatory T cell responses, its absence allows for the maintenance of Treg-mediated immune regulation.

### Aging-Associated Microbiota Induces Oxidative Stress and DNA Damage in Regulatory T Cells

To understand the mechanisms by which the microbiota of aged mice impairs Treg cell function, we performed pathway enrichment and gene expression analyses of Treg cells isolated from young and aged SPF and GF mice.

KEGG pathway enrichment analysis revealed that the Treg cells from old SPF mice upregulated genes related to pathways linked to oxidative phosphorylation, reactive oxygen species (ROS) production, apoptosis, TNF signaling, and p53 signaling **(Fig. 6A)**. In line with these findings, transcriptomic data reanalyzed from the study of Guo et al., (2020), showed that aged Treg cells exhibit increased expression of genes related to cellular stress, including Cdkn2a (p16) and Cdkn1a (p21), while genes involved in DNA repair, such as Rad9b and Xpc, were downregulated **(Fig. 6B)**. This shift in gene expression suggests that aging promotes a stress-prone transcriptional profile in Treg cells, potentially impairing their function and stability.

**Figure 6:**
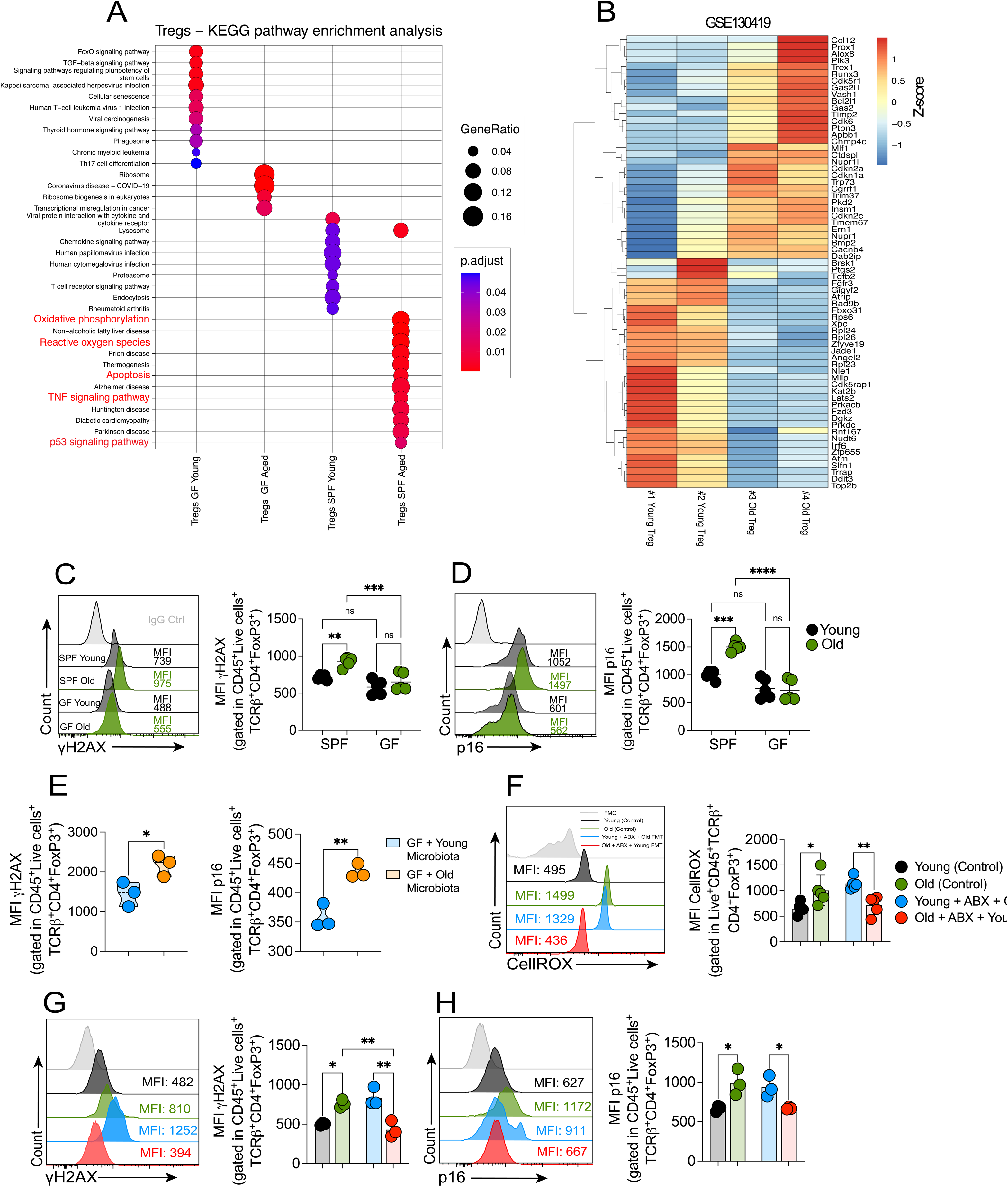
Aging-associated microbiota induces oxidative stress, DNA damage, and transcriptional dysregulation in Treg cells. (A) KEGG pathway enrichment analysis of differentially expressed genes in aged SPF Treg cells compared to young SPF Treg cells, highlighting enrichment for pathways related to oxidative phosphorylation, reactive oxygen species (ROS), apoptosis, TNF signaling, and p53 signaling. (B) Heatmap showing expression profiles of DNA damage response (DDR) genes in Treg cells from SPF-aged, SPF-young, GF-aged, and GF-young mice. Data were obtained from reanalyzed RNA-seq dataset GSE130419. Expression values are Z-score normalized. (C) Representative histograms and quantification of γH2AX expression (MFI) in Treg cells from SPF-aged, SPF-young, GF-aged, and GF-young mice. (D) Representative histograms and quantification of p16 expression (MFI) in Treg cells across the same groups as panel C. (E) Representative histograms and quantification of ROS levels in Treg cells from SPF-aged, SPF-young, GF-aged, and GF-young mice. ROS levels are measured by a fluorescent ROS probe. Data represent mean ± SEM. (G) Representative histograms and quantification of γH2AX expression (MFI) in Foxp3⁺ Treg cells from mice subjected to ABX treatment + FMT (aged control, young + aged FMT, aged + young FMT). Data shown as mean ± SEM. (H) Representative histograms and quantification of p16 expression (MFI) in Foxp3⁺ Treg cells from FMT-treated mice (same groups as in G). Data represent mean ± SEM. Data shown are representative of two to three independent experiments (n = 3–5 mice per group) and presented as mean ± SEM. Statistical analysis was performed using ANOVA with post-hoc test or unpaired t-test. *p < 0.05; **p < 0.01. Each symbol represents one mouse.

To directly evaluate whether this transcriptional reprogramming is associated with increased DNA damage, we measured γH2AX expression as a marker of DNA damage accumulation **(Fig. 6C)**. Aged SPF Treg cells exhibited significantly higher γH2AX levels compared to Treg cells from young SPF and from aged GF mice, indicating genomic instability. Notably, germ-free (GF) mice colonized with aged microbiota also displayed elevated γH2AX expression in Treg cells **(Fig. 6E)**. In parallel, we examined p16 expression, a marker linked to impaired proliferation and cellular stress responses. Aged Treg cells exhibited higher p16 expression **(Fig. 6D)**, and this phenotype was reduced in young mice that received aged microbiota **(Fig. 6H)**, further supporting the idea that microbiota composition plays a role in driving these stress-related changes.

Since oxidative stress is a key driver of both DNA damage and cellular dysfunction, we next assessed ROS levels in Treg cells using CellROX staining **(Fig. 6F)**. Aged Treg cells displayed significantly higher oxidative stress levels, and this phenotype was further exacerbated in young mice colonized with aged microbiota. These results suggest that microbiota composition influences oxidative stress in Treg cells, potentially leading to genomic instability and functional decline.

As these data indicate that aged Treg cells exhibit increased oxidative stress, DNA damage, and impaired proliferative capacity, we next sought to determine whether aging affects their ability to regulate these stress responses. To address this, we analyzed single-cell transcriptomic data to identify transcriptional regulators that might be altered in aged Treg cells. Among the differentially expressed transcription factors, c-Maf was significantly downregulated in SPF-aged Treg cells **(Supplementary** Fig. 1C), suggesting a potential role for this factor in maintaining Treg cell homeostasis under aging conditions. Given that c-Maf is involved in Treg cell stability and function^34–36^, we hypothesized that its downregulation could contribute to the inability of aged Treg cells to counteract cellular stress. Interestingly, we found a significant reduction in c-Maf expression in aged Treg cells compared to their young counterparts, suggesting that aging negatively impacts c-Maf expression. Notably, young FMT restored c-Maf expression in aged Treg cells, which was associated with decreased expression of p16 when compared to young mice that received old FMT **(Supplementary** Fig. 1D, E, F, G). To further investigate the role of c-Maf in Treg cell stability and stress responses, we analyzed young Foxp3^cre^Maf^flox/flox^ mice, which lack c-Maf specifically in Treg cells. Remarkably, γH2AX and p16 levels were significantly increased in c-Maf-deficient Treg cells compared to control Treg cells **(Supplementary Fig. Fig. 1H)**, indicating that the loss of c-Maf enhances susceptibility to DNA damage and cellular stress.

Together, these findings suggest that microbiota alterations associated with aging contribute to oxidative stress and DNA damage in Treg cells, potentially impairing their stability and function.

### Aging-Associated Microbiota Promotes Treg Cell Dysfunction Through Increased TNF Signaling

Our previous findings showed that aged Treg cells exhibit increased oxidative stress, DNA damage, and reduced proliferative capacity, impairments that were reversed by microbiota transfer from young donors (Fig. 6). However, the specific molecular mechanisms linking aging-associated microbiota to Treg dysfunction remained unclear. Given that KEGG pathway enrichment analysis identified TNF signaling as one of the most upregulated pathways in aged Treg cells (Fig. 6A), we hypothesized that chronic exposure to TNF, driven by microbial alterations during aging, contributes to Treg instability and functional decline.

To determine whether aged Treg cells are more responsive to TNF signaling, we evaluated TNFR1 and TNFR2 expression. Aged Treg cells displayed significantly higher TNFR1 expression in the mLNs and LI-LP compared to young Treg cells, while TNFR2 levels remained unchanged **(Fig. 7A)**. This suggests that aged Treg cells may be preferentially sensitized to TNF-driven inflammatory signals in the gut environment.

**Figure 7:**
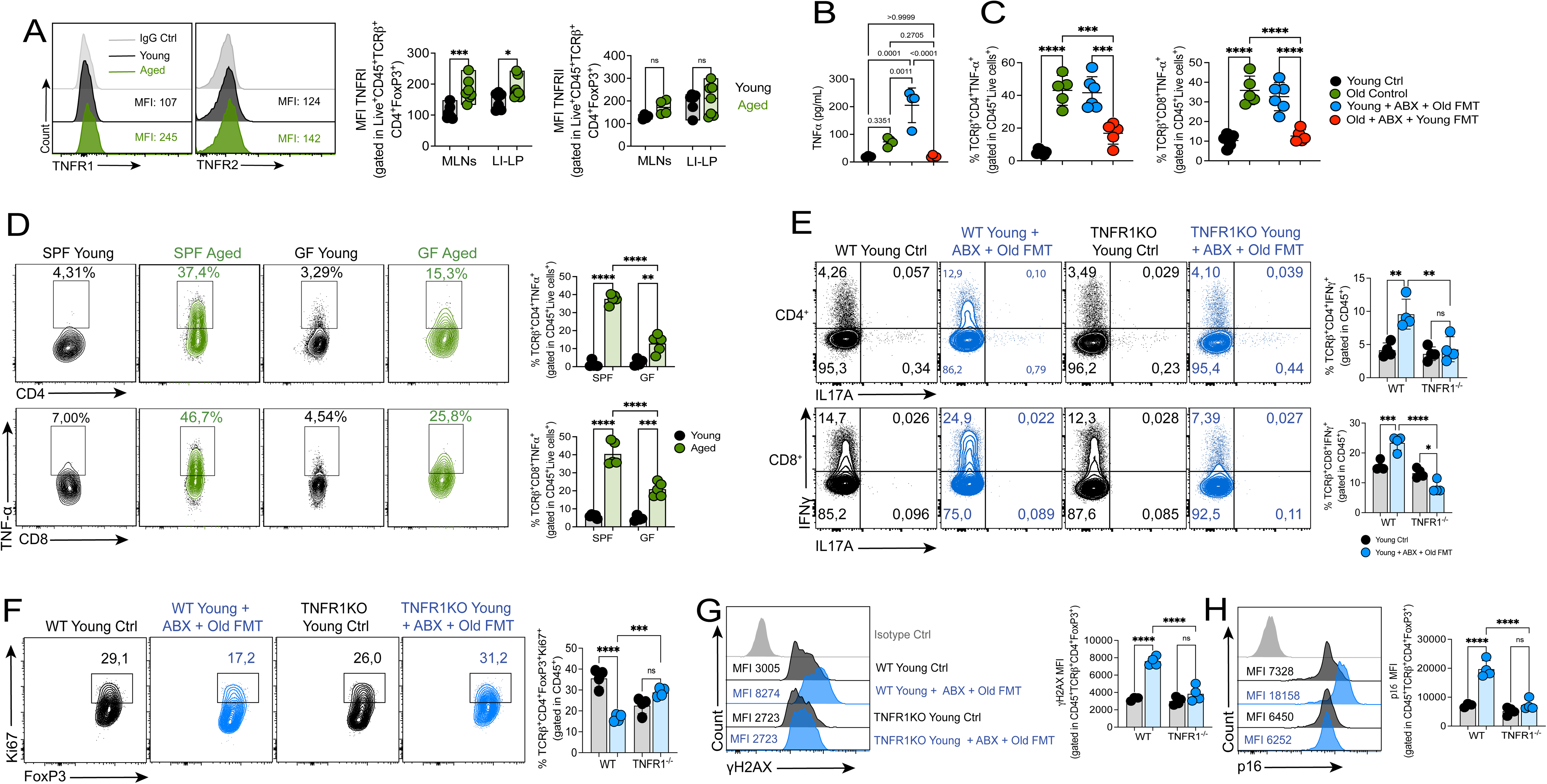
TNF–TNFR1 signaling mediates inflammatory T cell responses and Treg dysfunction driven by aged microbiota. (A) Representative histograms (LI-LP) and quantification of TNFR1 and TNFR2 expression (MFI) in Foxp3⁺ Treg cells from mesenteric lymph nodes (mLNs) and large intestinal lamina propria (LI-LP) of young and aged mice. (B) TNF levels (pg/mg) in colon tissue lysates from young, aged, antibiotic (ABX)-treated mice, and FMT recipients, measured by ELISA. (C) Quantification of TNF⁺ CD4⁺, and CD8⁺ T cells in the spleen of young, aged, ABX-treated, and FMT-recipient mice. (D) Representative dot plots and quantification of TNF⁺ CD4⁺, and CD8⁺ T cells in the spleen of SPF-aged and GF-aged mice. (E) Representative dot plots and quantification of IFN-γ⁺ and IL-17A⁺ CD4⁺ and CD8⁺ T cells in the spleen of WT control, WT + ABX + aged FMT, TNFR1KO control, and TNFR1KO + ABX + aged FMT mice. (F) Quantification of Ki67⁺ proliferating Foxp3⁺ Treg cells in WT control, WT + ABX + aged FMT, TNFR1KO control, and TNFR1KO + ABX + aged FMT mice. (G) Representative histograms and quantification of γH2AX expression (MFI) in Foxp3⁺ Treg cells from the same groups as in panel F. (H) Representative histograms and quantification of p16 expression (MFI) in Foxp3⁺ Treg cells from WT and TNFR1KO mice under control and ABX + aged FMT conditions. Data shown are representative of two to three independent experiments (n = 4–8 mice per group) and presented as mean ± SEM. Statistical analysis was performed using ANOVA with post-hoc test or unpaired t-test. *p < 0.05; **p < 0.01. Each symbol represents one mouse.

To further investigate the role of microbiota in TNF signaling, we first measured TNF levels in colonic tissue following FMT **(Fig. 7B)**. Aged SPF mice exhibited significantly higher TNF levels compared to young SPF controls, confirming an aging-associated increase in TNF production. Importantly, young FMT into aged mice significantly reduced colonic TNF levels, whereas transferring aged microbiota into young mice led to an increase in TNF levels **(Fig. 7B)**. These findings suggest that microbiota composition is a key regulator of TNF production in the aging gut.

Next, we assessed TNF production at the cellular level using flow cytometry to quantify TNF-expressing CD4⁺ and CD8⁺ T cells in the colon after FMT. Consistent with the ELISA results, aged SPF mice displayed a significant increase in the percentage of TNF⁺ CD4⁺ and CD8⁺ T cells compared to young SPF controls **(Fig. 7C)**. Of note, young FMT into aged mice reduced TNF production in both CD4⁺ and CD8⁺ T cells, while aged microbiota transfer into young mice led to a marked increase in TNF⁺ T cells **(Fig. 7C)**.

These data further support the idea that aging-associated microbiota actively contributes to TNF-driven immune activation in the gut.

To determine whether this microbiota-driven TNF production is dependent on the aged-microbiota, we analyzed TNF expression in both young and old SPF and GF mice **(Fig. 7D)**. Aged SPF mice exhibited a pronounced increase in TNF⁺ CD4⁺ and CD8⁺ T cells. In contrast, aged GF mice displayed significantly lower frequencies of TNF⁺ T cells, indicating that microbiota-derived signals fuel TNF-driven immune activation. These results collectively demonstrate that the aging microbiota promotes a TNF-enriched inflammatory environment, which may contribute to Treg cell dysfunction and immune dysregulation during aging.

To directly assess whether TNF signaling via TNFR1 contributes to the immune alterations induced by aged microbiota, we performed FMT from aged WT donors into young WT or TNFR1-deficient (TNFR1KO) mice pre-treated with antibiotics. As expected, WT recipients of aged microbiota developed an increased frequency of IFN-γ⁺ CD4⁺ and CD8⁺ T cells in the colon, consistent with an inflammatory environment **(Fig. 7E)**. In contrast, TNFR1KO mice receiving the same FMT exhibited significantly lower frequencies of these effector subsets, suggesting that the absence of TNFR1 reduces susceptibility to microbiota-induced immune activation. We then examined the phenotype of Foxp3⁺ Treg cells in these animals. Compared to WT mice, TNFR1KO recipients displayed higher proportions of Ki67^+^ Treg cell **(Fig. 7F)**. Furthermore, expression of the DNA damage marker γH2AX and the senescence-associated protein p16 was significantly lower in Treg cell from TNFR1KO mice colonized with aged microbiota **(Fig. 7G-H)**, suggesting reduced cellular stress and improved homeostasis in the absence of TNFR1 signaling.

Together, these findings support the idea that TNF–TNFR1 signaling contributes to the immunological alterations associated with aged microbiota, including effector T cell expansion and Treg dysfunction.

### Aging Induces Transcriptional Remodeling in Human Splenic Treg Cells

To investigate how aging impacts the transcriptional landscape of human Treg cells, we analyzed the publicly available single-cell multimodal dataset from Wells et al. (2024)^37^, which profiled immune cells from multiple tissues of organ donors aged between 20 and 75 years using CITE-seq. For this analysis, we focused on splenic Treg cells, as the spleen was the tissue with the highest number of Treg cells recovered, providing greater resolution for transcriptional comparisons.

Differential gene expression analysis comparing splenic Treg cell from young (<40 years) and aged (>=40 years) donors, while controlling for the influence of donor CMV status, sex, sample collection site and assay employed, revealed a substantial transcriptional shift with aging. Aged Treg cells displayed consistently higher expression of a broad set of genes involved in TNF signaling, including NFKBIA, NFKBIZ, TNFAIP3, JUNB, DDIT4, CXCR4, KLF6, PPP1R15A, and HSPA5, which were uniformly upregulated in the aged group (**Fig. 8A**).

**Figure 8:**
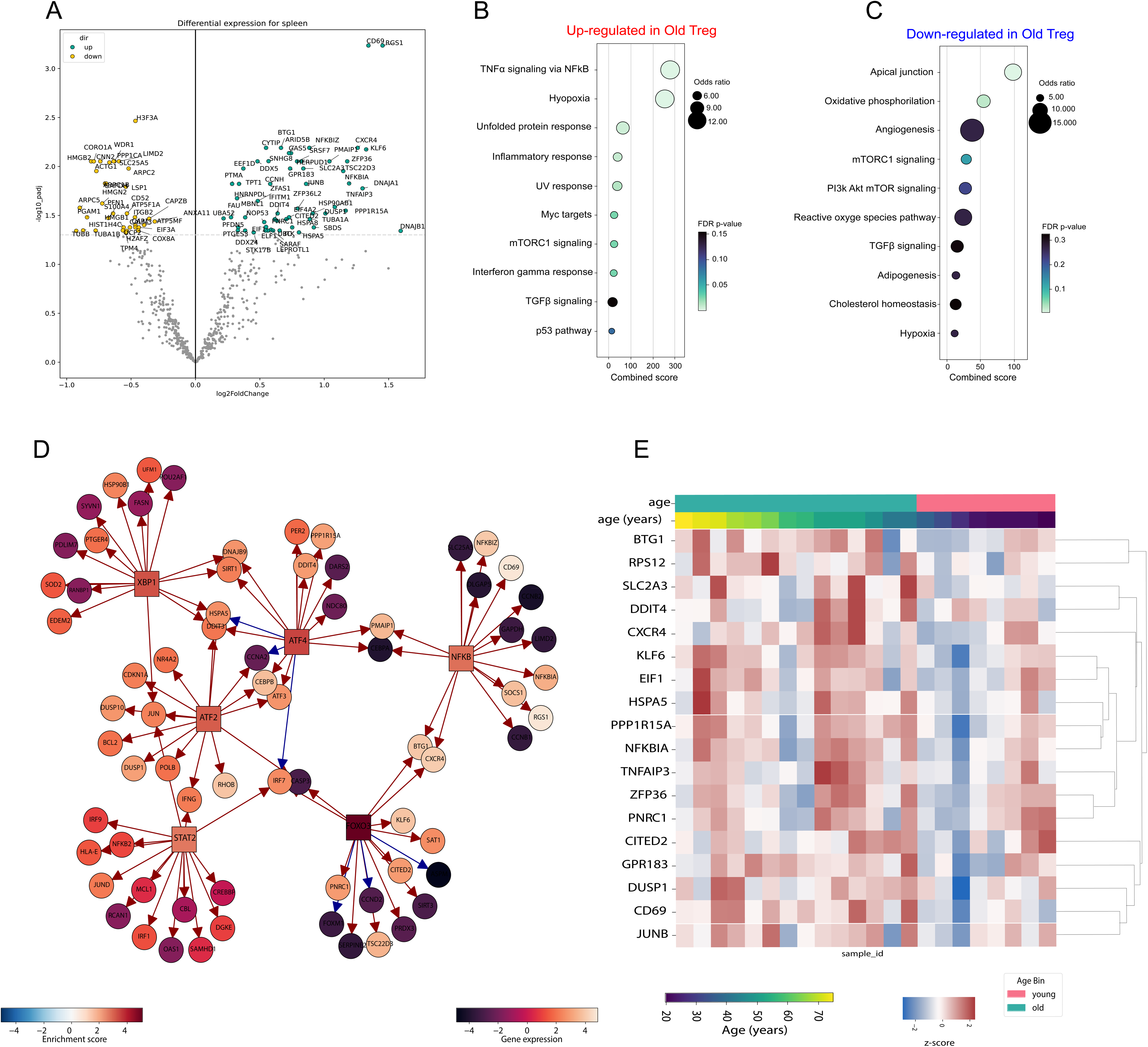
Transcriptional and regulatory landscape of splenic Treg cells from young and aged humans. (A) Volcano plot showing differentially expressed genes (DEGs) in Treg cells isolated from human spleens comparing young (<40 years) and aged (>=40 years) donors. Differential expression analysis was performed on aggregated pseudobulk data and consecutive DESeq2, with correction for multiple biological and technical confounders (sex, CMV status, employed assay and sample processing site). Highlighted genes are significantly upregulated (orange) or downregulated (blue) in aged Treg cells. (B-C) Over-representation analysis (ORA) of pathways enriched among genes upregulated (B) and downregulated (C) in aged Treg cells relative to young controls, based on DEG results shown in (A). (D) Transcription factor (TF) regulatory network inferred from the DEG set, showing TFs (squares) and their downstream target genes (circles). The TF node color indicates whether the TF is enriched in aged (red) or young (blue) Treg cells. The color of the gene nodes represents the scaled gene expression across the Treg population. Arrows denote positive regulation, while bars indicate negative regulatory effects on downstream targets. (E) Heatmap of scaled expression values for the DEGs between young and aged splenic Treg cells. Columns represent individual donors stratified by age, and rows represent genes.

Pathway enrichment analysis revealed that the genes upregulated in aged Treg cells were associated with cellular stress responses, inflammatory signaling, and cytokine-mediated pathways. The most enriched terms included unfolded protein response, TNF signaling, NF-κB signaling, and cellular response to stress (**Fig. 8B**). Conversely, the genes downregulated in aged Treg cells showed enrichment for pathways related to oxidative phosphorylation and TGF® signaling (**Fig. 8C**).

To further dissect the regulatory architecture underlying these transcriptional changes, we performed transcription factor network inference. This analysis revealed increased activity of NF-κB (NFKB1/NFKBIA), STAT1, IRF1, JUN, FOS, ATF4, and HIF1A in aged Treg cells, driving regulatory circuits linked to TNF signaling, inflammation, and stress responses. In addition, the network includes key senescence– and stress-associated genes such as DDIT4, PPP1R15A, KLF6, and HSPA5, all consistently upregulated in aged Treg cells. Pro-inflammatory targets like NFKBIA, TNFAIP3, CD69, CITED2, CXCR4, and GPR183 further characterize this aged-associated signature (**Fig. 8D**).

These transcriptional differences are clearly reflected in the heatmap, which demonstrates a robust and consistent segregation of donors according to age. Aged Treg cells are characterized by uniformly elevated expression of genes analyzed enriched for TNF signaling, NF-κB signaling, and cellular response to stress, including NFKBIA, TNFAIP3, JUNB, DDIT4, CXCR4, KLF6, PPP1R15A, HSPA5, DUSP1, CD69, CITED2, GPR183, RPS12, EIF1, and SLC2A3 (**Fig. 8E**)

Collectively, these data reveal that aging drives a transcriptional reprogramming in human splenic Treg cells, characterized by a coordinated upregulation of genes involved in stress responses, inflammation, and activation, alongside the suppression of a limited set of genes associated with proliferative capacity and homeostatic functions which goes in line with our findings in mouse aged Treg.

## Discussion

Aging is marked by shifts in immune homeostasis, characterized by low-grade inflammation and impaired regulation. Here, we identify a previously unrecognized axis through which microbiota-driven inflammatory signaling and cellular stress responses converge to destabilize regulatory T cell function with age. We show that aged mice accumulate Treg cells with an effector-memory phenotype, diminished proliferative capacity, and selective changes of suppressive pathways, impairing their ability to restrain pro-inflammatory T cell responses. A central finding of this work is that microbial dysbiosis associated with aging is sufficient to drive this Treg dysfunction by promoting oxidative stress and DNA damage through TNF–TNFR1 signaling, compromising Treg stability and function.

Previous studies had indicated that Treg numbers expanded with age ^7,10,14,16^, possibly in a compensatory manner to increased immune stimulation. However, how aging affects the function of these cells is unclear. Our results complement these findings by demonstrating that aged Treg cells, although numerically expanded, have functional defects such as diminished proliferation, enhanced terminal differentiation (KLRG1⁺), and an inability to suppress T cell-mediated colitis. Importantly, the increased frequency of Tregs in aged mice does not reflect enhanced turnover or homeostatic proliferation, as evidenced by reduced expression of Ki67, a marker of active cycling. Instead, this accumulation may result from impaired apoptosis or preferential survival of senescent or dysfunctional Tregs. Supporting this, studies have shown that aged Tregs exhibit features of cellular senescence, including elevated expression of p16^INK4a and DNA damage markers such as γH2AX ^10,18^.

The increased expression of KLRG1 observed in aged Tregs in our study aligns with recent findings identifying KLRG1 as a marker of regulatory T cell (Treg) subsets that accumulate with age and exhibit features of cellular senescence. A recent study by Soto-Heredero et al. (2025) demonstrated that KLRG1⁺ Tregs (kTregs) progressively expand in aged mice and humans, and display hallmark features of senescence, including mitochondrial dysfunction, elevated expression of p16^INK4a, and increased DNA damage markers such as γH2A X. Although these cells retained suppressive capacity in vitro, their function was impaired in vivo and accompanied by a pro-inflammatory phenotype, suggesting functional fragility of aged kTregs ^18^. Additionally, KLRG1 signaling has been shown to directly impair T cell proliferation through inhibition of AKT phosphorylation and induction of cell cycle inhibitors such as p16 and p21, reinforcing its role as an active regulator of senescence and dysfunction in both CD4⁺ and CD8⁺ T cells^38,39^. These findings offer a mechanistic explanation for our observation that aged Tregs, despite their numerical expansion, exhibit reduced proliferative capacity (low Ki67 expression) and a terminally differentiated, dysfunctional phenotype. Thus, KLRG1 expression may not only mark but also contribute to Treg instability and impaired immune regulation during aging.

One of the key findings of our study is the causal role of the gut microbiota in this process. In fecal microbiota transplantation and cohousing experiments, we demonstrate that exposure to aged microbiota alone is sufficient to impair Treg proliferative capacity and generate a pro-inflammatory T cell environment even in young hosts. Reconstitution with a young microbiota, however, restores Treg function in old mice. In addition to these findings, our results in GF mice provide clear evidence that the microbiota is essential for driving the immune alterations seen with aging. Aged GF mice retain Treg cells with preserved proliferative capacity and lower pro-inflammatory T cell frequencies, contrasting with aged SPF mice, where Treg cells become dysfunctional. Remarkably, transferring aged microbiota into young GF mice is sufficient to induce the same inflammatory skewing and Treg impairment, confirming the microbiota as a key driver of immunosenescence. These results align with previous studies showing that aged microbiota triggers systemic inflammation, enhances leakage of microbial products (TLR2/TLR4 ligands), and upregulates TNF-α as a central mediator of inflammatory gene expression ^22^. Additionally, aged microbiota reduces short-chain fatty acids (SCFAs), key metabolites that support Treg stability, while enriching inflammatory taxa like Proteobacteria and depleting beneficial SCFA producers ^40^. Therefore, the altered Treg function in the elderly could also be due to decreased SCFA accessibility.

Mechanistically, we demonstrate that oxidative stress and DNA damage are key mediators of Treg cell dysfunction in aged SPF mice. Treg cells from these animals exhibit elevated ROS levels, increased γH2AX accumulation (a marker of DNA double-strand breaks), and p16 upregulation, features characteristic of cellular senescence. Notably, these stress signatures are significantly reduced in aged GF mice, reinforcing that microbial cues are necessary to induce genomic instability in Treg cells. This is consistent with evidence that DNA damage is a central driver of the aging process, contributing to genomic instability, epigenetic alterations, mitochondrial dysfunction, and stem cell exhaustion ^41^. In particular, the persistence of DNA lesions and the resulting DNA damage response (DDR) promote senescence and inflammatory signaling, which are hallmarks of aging tissues ^42,43^. Our findings are reflected on this, as Treg cells from aged SPF mice accumulate γH2AX and express p16, indicative of chronic DDR activation, while Treg cells from aged GF mice maintain genomic integrity and suppressive potential. This suggests that microbiota-derived signals, possibly through inflammatory mediators, accelerate Treg cell senescence by enhancing oxidative stress and DNA damage.

Moreover, we identified c-Maf as a critical regulator linking microbial signals, inflammatory stress, and Treg cell stability. c-Maf expression is significantly reduced in Treg cells from aged SPF mice, suggesting that microbiota-driven inflammation suppresses c-Maf levels. Additionally, c-Maf-deficient Treg cells accumulate higher levels of γH2AX and p16, indicative of increased DNA damage and cellular senescence. This suggests that c-Maf could play a role in protecting Treg cells from oxidative and genotoxic stress, likely by modulating antioxidant responses and maintaining chromatin homeostasis. However, although c-Maf has been previously implicated in the regulation of Treg function and stability ^34–36^, it remains unclear whether c-Maf loss in aged animals fully explains the extent of Treg cell dysfunction observed during aging. Due to the lack of aged c-Maf-deficient mice, we could not directly test whether aging synergizes with c-Maf deficiency to further exacerbate Treg instability. Future studies using c-Maf-deficient mice at advanced ages will be necessary to understand how c-Maf modulates DNA damage responses and Treg cell homeostasis in the context of aging and chronic inflammation.

In our study, we reveal that chronic TNF production in aged mice shaped by microbiota dysbiosis directly contributes to genomic instability in Treg cells, alongside impaired proliferative fitness. In this sense, a previous study demonstrated that the aged microbiota promotes intestinal barrier disruption and elevates systemic TNF levels, positioning TNF as a central mediator of age-associated inflammation and microbial dysbiosis ^44^. In our model, this inflammatory axis translates into increased DNA damage and reduced proliferation in Treg cells. While TNF-α is widely recognized as a hallmark cytokine of inflammaging, our data extend this concept by revealing that aged Treg cells, which upregulate TNFR1, are particularly vulnerable to TNF-induced genotoxic stress.. In line with these, a previous study has shown that TNF-α is sufficient to induce DNA damage across multiple cell types, including T cells, both in inflammatory models and upon systemic TNF exposure ^45^, our data suggest that in Treg cells, ROS accumulation likely plays a dominant role, converging with TNFR1 signaling to destabilize genomic integrity. Although systemic TNFR1 deficiency ameliorates these defects, the precise contribution of TNFR1 signaling within Treg cells remains unresolved. Whether intrinsic TNFR1 activation directly triggers DNA damage responses and senescence pathways in Treg cells during aging, or whether this reflects the broader inflammatory environment, remains an open question. Addressing this will require Treg cell-specific TNFR1-deficient models at advanced ages, which could define TNFR1 as a cell-intrinsic regulator of Treg cell genomic stability. Supporting this notion, our transcriptional analysis of human splenic Treg cells reveals a conserved signature of aging characterized by chronic activation of TNF– and stress-related pathways. Aged human Treg cells show increased activity of key transcription factors, including NF-κB, STAT1, IRF1, and ATF4, coupled with robust upregulation of genes associated with cellular stress, senescence, and proteostasis dysfunction, such as DDIT4, PPP1R15A, KLF6, and HSPA5. Notably, several of these genes are downstream targets of TNF signaling and are tightly linked to senescence-associated transcriptional programs in other systems. These findings suggest that the TNFR1-driven stress axis identified in mice is similarly engaged in human Treg cells during aging.

Our findings offer conceptual insights that may inform therapeutic strategies aimed at restoring immune balance in aging. Given the central role of chronic inflammation and immune dysregulation in age-related diseases—including inflammatory bowel disorders, metabolic dysfunction, and cancer—intervening along the microbiota–TNF–Treg cell axis could represent a promising avenue. Approaches that reshape the microbial landscape, modulate TNF–TNFR1 signaling, or enhance Treg cell resilience, for example by sustaining c-Maf expression, may help preserve Treg cell stability and genomic integrity, thereby mitigating the progressive loss of immune regulation that accompanies aging.

In summary, our studies show that microbial dysbiosis associated with aging induces Treg cell dysfunction through TNF–TNFR1-mediated oxidative stress and DNA damage, and disrupts immune homeostasis and inflammation associated with aging. These findings present a mechanistic paradigm for the interaction between microbiota, inflammatory signaling, and Treg cell homeostasis and indicate potential therapeutic targets for immune aging modulation.

## Materials and Methods

### Mice

C57BL/6J mice were maintained under either specific pathogen-free (SPF) or germ-free (GF) housing conditions. GF mice were generated by embryo transfer and housed in sterile flexible-film isolators^46^. GF status was confirmed weekly by PCR detection of 16S rDNA and bacterial cultures. Young mice were 4–12 weeks old, and aged mice were 70 weeks old. All procedures performed on mice were approved by the local committee on legislation on protection of animals (Landesuntersuchungsamt Rheinland-Pfalz, Koblenz, Germany; 00033240-1-X)

### Fecal Microbiota Transfer (FMT) and Antibiotic Treatment

For microbiota depletion, recipient mice were treated daily via gavage with ampicillin (1.86 mg), vancomycin (0.96 mg), neomycin sulfate (1.86 mg), and metronidazole (1.86 mg) dissolved in 300 μl of sterile water for 2 weeks. After depletion, fecal material was freshly collected from young (4–12 weeks) or aged (70 weeks) SPF donors, diluted (100 mg/mL) in sterile PBS under anaerobic conditions, homogenized, filtered, and administered by oral gavage (300 μl per mouse) for 5 consecutive days. For GF mice, the same protocol was applied without prior antibiotic treatment.

### Histology and Hematoxylin & Eosin (H&E) Staining

Segments of distal colon were harvested, flushed with cold PBS to remove fecal contents, and fixed in 4% paraformaldehyde (PFA) at 4°C for 24 hours. Tissues were then dehydrated through a graded ethanol series, cleared in xylene, and embedded in paraffin. Sections of 5 µm thickness were cut using a microtome and mounted on glass slides. After deparaffinization and rehydration, sections were stained with hematoxylin for nuclear visualization, followed by eosin for cytoplasmic and extracellular matrix contrast. Slides were dehydrated, cleared, and mounted with coverslips using a permanent mounting medium. Images were acquired using a light microscope (e.g., Leica or Olympus) under 10x and 20x magnifications.

### In vivo high-resolution mini-endoscopy analysis of the colon

To monitor colitis progression in live animals, high-resolution mini-endoscopy was performed using a veterinary endoscopic system (Karl Storz, Germany) following the protocol previously described by Reißig et al. (2017)^47^. Mice were anesthetized via intraperitoneal injection of ketamine (Ketavest, 100 mg/ml; Pfizer) and xylazine (Rompun, 2%; Bayer Healthcare). Following anesthesia induction, a rigid endoscope was gently inserted into the rectum, and video recordings of the distal colon were obtained at designated time points. Colitis severity was assessed using a validated endoscopic scoring system based on five parameters: mucosal translucency, granularity, presence of fibrin, vascular pattern, and stool consistency. Each parameter was scored individually, and the total score was used to quantify colonic inflammation. All procedures were performed under sterile conditions, and scoring was conducted in a blinded manner.

### In vitro Treg Suppression Assay

The suppressive capacity of regulatory T cells (Tregs) from young and aged mice was evaluated using a standard in vitro suppression assay, as previously described by Collison and Vignali^48^ with modifications. Tregs were isolated from spleens and mesenteric lymph nodes of young or aged mice (maintained either in single housing or cohoused with young mice for four weeks) using a CD4⁺CD25⁺ Regulatory T Cell Isolation Kit (Miltenyi Biotec) following the manufacturer’s instructions. Naive CD4⁺ T cells (CD4⁺CD62L⁺CD25⁻) from young mice were isolated using the Naive CD4⁺ T Cell Isolation Kit (Miltenyi Biotec) and subsequently labeled with CellTrace™ Violet (CTV, ThermoFisher) according to the manufacturer’s protocol. CTV-labeled naive CD4⁺ T cells were cultured either alone or in the presence of Tregs at a 1:1 ratio (1×10⁵ cells each) in 96-well round-bottom plates pre-coated with anti-CD3 (2 µg/mL, clone 145-2C11) and soluble anti-CD28 (1 µg/mL, clone 37.51). Cells were cultured in complete RPMI 1640 medium supplemented with 10% heat-inactivated fetal bovine serum, 1% penicillin-streptomycin, 2 mM L-glutamine, 1 mM sodium pyruvate, non-essential amino acids, 10 mM HEPES, and 50 µM 2-mercaptoethanol. Cultures were maintained at 37°C and 5% CO₂ for 72 hours. After incubation, cell proliferation was assessed by flow cytometric analysis of CTV dilution in the CD4⁺ Tconv population.

### Co-Housing Experiments

To assess the impact of microbiota transfer via co-housing, female C57BL/6J mice were used. Aged (70 weeks) and young (4–12 weeks) mice were co-housed at a 1:1 ratio in standard cages under specific pathogen-free (SPF) conditions for 4 weeks. Control groups of aged and young mice were maintained in single housing under the same conditions. Food, water, and bedding were changed simultaneously across all cages to minimize environmental variation. At the end of the co-housing period, mice were euthanized, and tissues including spleen, mesenteric lymph nodes (mLNs), and intestinal lamina propria (cLP and SILP) were collected for downstream flow cytometry, functional assays, and microbiota analysis.

### Isolation of Lamina Propria Lymphocytes

Lamina propria lymphocytes from the colon (cLP) and small intestine (SILP) were isolated following standard protocols ^49,50^. Briefly, the intestines were harvested, cleaned, and epithelial cells were removed via EDTA/DTT washes. Tissue was digested using collagenase VIII (0.5 mg/mL) or Liberase (Roche) (5 mg/mL) and DNase I (0.5 mg/mL) at 37°C for 25 min under agitation. The resulting cell suspensions were filtered for debris removal and washed for downstream assays.

### Flow Cytometry

Phenotypic and functional analysis of lymphocyte populations isolated from the spleen, mesenteric lymph nodes (mLNs), visceral adipose tissue (VAT), lung and intestinal lamina propria (cLP and SILP) was performed by flow cytometry. Following isolation, cells were counted and assessed for viability using trypan blue exclusion. For staining, 2 × 10⁶ cells per well were plated in 96-well plates. Cells were first incubated with Fc receptor blockade to prevent non-specific antibody binding. After washing, surface staining was performed using antibodies against TCRb, CD45, CD8, CD4, CD44, CD62L, CTLA-4, CD39, ICOS, PD-1 and TNFR1, along with a fixable viability dye. Subsequently, cells were fixed and permeabilized using a transcription factor staining buffer set. Intracellular staining included markers for IFNg, IL-17, Foxp3, RORgt, Helios, Ki67, γH2AX, p16, and c-Maf, performed at 4°C in permeabilization buffer. For reactive oxygen species (ROS) detection, cells were incubated with a ROS-sensitive fluorescent probe prior to fixation, following the manufacturer’s guidelines. Data acquisition was carried out on a BD LSR Fortessa flow cytometer, and analysis was performed using FlowJo software. Compensation controls and fluorescence minus one (FMO) control were employed to define gating strategies.

### *In Vivo* Treg Cell Suppression Assay

To assess Treg suppressive function *in vivo*, MACS-purified naïve CD4⁺CD45RB^hi^ T cells (5 × 10⁵) were co-transferred with Treg cells (5 × 10⁵) isolated from young or aged donors into 6-8-week old Rag1⁻/⁻ mice via intravenous injection. Body weight was monitored weekly as a marker of disease progression. After 4–6 weeks, colon length and inflammatory cytokine production were assessed.

### 16S rRNA Gene Sequencing and Microbiota Analysis

Microbiota profiling was performed by Novogene Europe. DNA was extracted from fecal samples and subjected to 16S rRNA gene amplicon sequencing targeting the V3–V4 regions on the Illumina MiSeq platform (PE250), yielding ∼30,000 reads per sample. Standard bioinformatics analysis, including OUT clustering, taxonomic annotation, and diversity metrics (alpha/beta-diversity), was conducted.

### Single Cell Analysis

Single-cell RNA sequencing (scRNA-seq) data were obtained from a previously published study, deposited in the DNA Data Bank of Japan (accession code: E-GEAD-583). Four datasets corresponding to distinct experimental groups (GF_Young, GF_Aged, SPF_Young, SPF_Aged) were imported using Read10X() and converted into Seurat objects via CreateSeuratObject() (min.cells = 3, min.features = 200). The datasets were merged for unified analysis. Quality control metrics were calculated using PercentageFeatureSet() to assess the proportion of mitochondrial (^mm10---mt-), ribosomal (^mm10---Rpl|^mm10---Rps), and globin (^mm10---Hba|^mm10---Hbb) transcripts. Cells with low complexity or signs of stress (nFeature_RNA < 200 or > 6000; nCount_RNA > 20,000; percent.mito > 12%; percent.globin > 0.1) were excluded. Data were normalized (NormalizeData()), and highly variable features were identified using the “vst” method. Scaling was performed with regression of ribosomal content (ScaleData(vars.to.regress = ‘percent.ribo’)). Dimensionality reduction was performed using PCA (RunPCA()), and the optimal number of principal components was selected using ElbowPlot(). Clustering was performed using the Louvain algorithm (FindNeighbors(), FindClusters(), resolution = 0.2), with visualization via UMAP and t-SNE embeddings. Doublets were detected and removed using the DoubletFinder package. After filtering, data were split by sample, re-normalized, and integrated using canonical correlation analysis (FindIntegrationAnchors(), IntegrateData()), followed by rescaling and dimensionality reduction. Gene module activity, including senescence-related signatures, was assessed using AddModuleScore(). The clusterProfiler package was used to perform Gene Ontology (GO) and KEGG pathway enrichment. Cluster-specific differentially expressed genes (DEGs) were grouped, and gene symbols were converted to Entrez IDs using the bitr() function. For each cluster, gene lists were assembled and passed to compareCluster() to conduct over-representation analysis for GO terms (enrichGO) and KEGG pathways (enrichKEGG). Enrichment results were visualized using dotplot() to highlight biologically enriched processes across clusters.

### ScRNA-Seq Analysis of Human Treg Cells

To investigate the impact of age on human Treg cells, we accessed CITE-seq data of Wells et al. (2024)^37^, which analyzed immune cells from multiple tissues of organ donors aged between 20 and 75 years. Splenic Treg cells were selected based on the original author’s annotations. To analyze the impact of age on Treg cells, we classified donors in two age groups based on a binary cutoff of the respective donor age (old >= 40; young <40). We then performed pseudobulk aggregation for Treg cells from each donor. Samples containing fewer than 10 Treg cells were excluded from the analysis. Differential gene expression between Treg cells of old and young donors was carried out using DESeq2^51^ while controlling for the influence of CMV status, sex, the site the data was collected and the assay employed to generate the data. Pathway enrichment analysis has been conducted using the MSigDB Hallmark collection and performing an overrepresentation analysis separately for upregulated and downregulated differentially expressed genes using decoupler^52^. Transcription factor inference has been performed using a univariate linear model in decoupler.

### Statistical Analysis

Statistical analyses were performed using GraphPad Prism 9. Data were tested for normality (Shapiro-Wilk test). Comparisons between two groups were performed using unpaired t-tests (parametric) or Mann-Whitney U tests (non-parametric). For multiple group comparisons, one-way ANOVA or Kruskal-Wallis tests were used, followed by appropriate post-hoc corrections. P-values <0.05 were considered statistically significant. Sample sizes (n) are indicated in each figure legend.

## Acknowledgments

We are grateful to Michaela Blanfeld and Elena Zurkowski for their technical assistance.

## Authors Contributions

J.A.L. designed and performed experiments, analyzed data, and wrote the manuscript. Z.E., E.S., R.J., N.N.B., A.P.Z., K.C., and X.L. assisted with experiments and data analysis. E.S, M.M; F.I. and N.Y. contributed to bioinformatic analyses. N.H., T.R., and C.N. provided technical support and critical discussion. C.R. contributed to experimental design and data interpretation. A.W. supervised the project and provided conceptual guidance. All authors reviewed and approved the final manuscript.

## Figures Legends

**Supplementary Figure 1:**
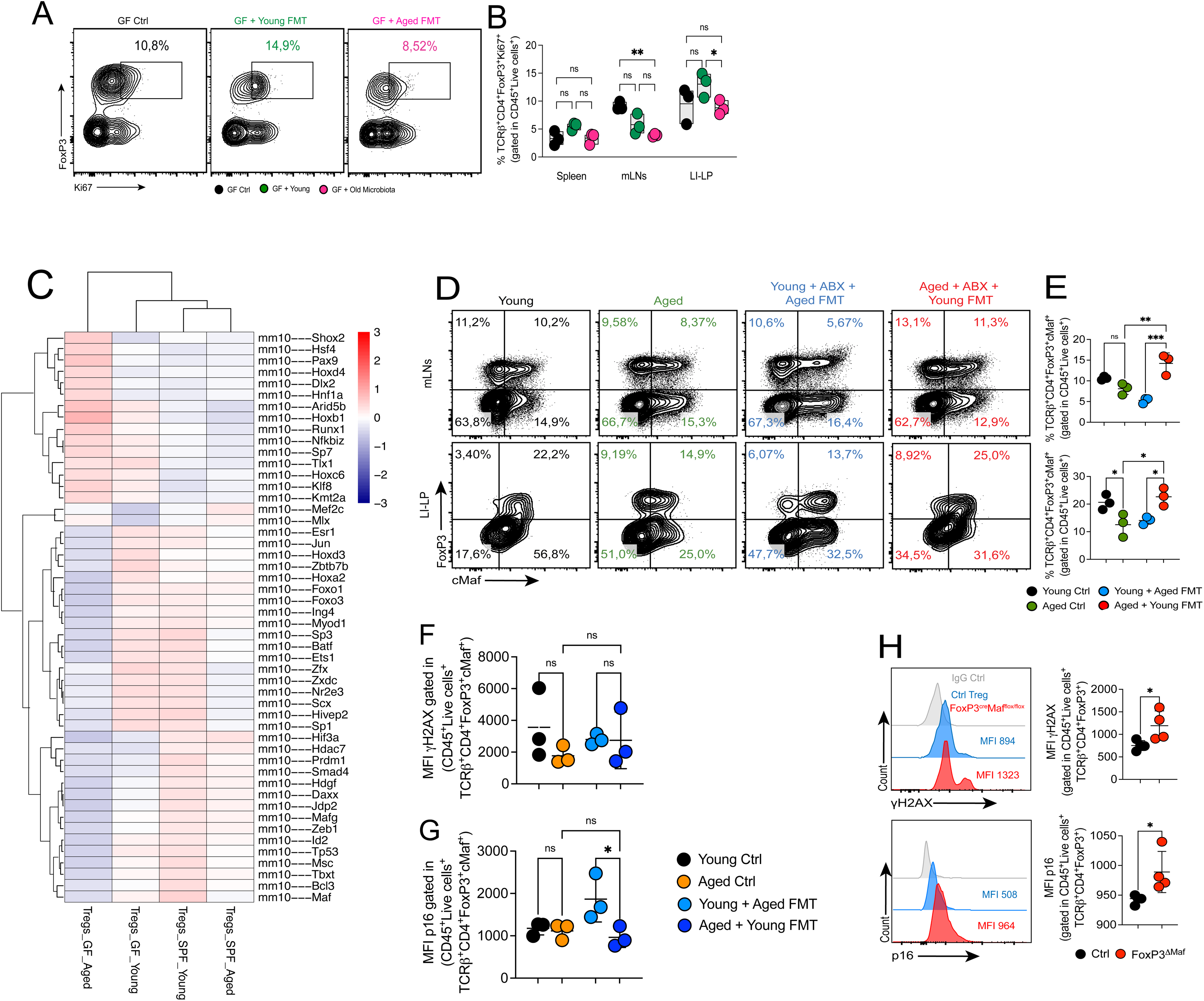
Microbiota transfer modulates Treg proliferation and inflammatory profile in germ-free mice. (A) Representative dot plots showing Foxp3⁺ Treg frequencies in the spleen of germ-free (GF) control mice, GF + young FMT, and GF + aged FMT recipients. The adjacent bar graph quantifies Treg cell frequencies (%). (B) Quantification of Ki67⁺ proliferating Treg cells in the mesenteric lymph nodes (mLNs) and large intestinal lamina propria (LI-LP) of GF control, GF + young FMT, and GF + aged FMT groups. (C) Heatmap showing Z-score normalized expression of TNF signaling-related genes in Treg cells from GF + young FMT and GF + aged FMT mice. Increased expression of inflammatory mediators is observed in aged FMT recipients. Color scale: red (high expression), blue (low expression). (D) Representative dot plots and quantification of c-Maf expression (%) in Foxp3⁺ Treg cells from FMT-treated mice (aged control, young + aged FMT, aged + young FMT). (E) Quantification of Foxp3⁺c-Maf⁺ Treg frequencies (%) and Foxp3⁺c-Maf⁻ Treg frequencies (%) in FMT-treated mice (aged control, young + aged FMT, aged + young FMT). (F, G) Quantification of γH2AX and p16 expression (MFI) in Foxp3⁺ Treg cells from young and aged control and after FMT. (H) Representative histograms and quantification of γH2AX and p16 expression (MFI) in Foxp3⁺ Treg cells isolated from the lamina propria (LP) of the colon from control mice and Foxp3^cre^Maf^flox/flox^ mice. Data shown are representative of two to three independent experiments (n = 4–5 mice per group) and presented as mean ± SEM. Statistical analysis was performed using ANOVA with post-hoc test or unpaired t-test. *p < 0.05; **p < 0.01. Each symbol represents one mouse.

